# Adaptations for bipedal walking: Musculoskeletal structure and three-dimensional joint mechanics of humans and bipedal chimpanzees (*Pan troglodytes*)

**DOI:** 10.1101/2022.02.21.481231

**Authors:** Matthew C. O’Neill, Brigitte Demes, Nathan E. Thompson, Susan G. Larson, Jack T. Stern, Brian R. Umberger

**Affiliations:** Department of Anatomy, Midwestern University, Glendale, AZ 85308 USA; Department of Anatomical Sciences, Stony Brook University School of Medicine, Stony Brook, NY 11794 USA; Department of Anatomy, New York Institute of Technology College of Osteopathic Medicine, Old Westbury, NY 11568 USA; School of Kinesiology, University of Michigan, Ann Arbor, MI 48109-2013 USA

**Keywords:** Force, Work, Power, Elastic Energy, Bipedalism, Locomotion

## Abstract

Humans are unique among apes and other primates in the musculoskeletal design of their lower back, pelvis and lower limbs. Here, we describe the three-dimensional ground reaction forces and lower/hind limb joint mechanics of human and bipedal chimpanzee walking over a full stride and test whether: 1) the estimated limb joint work and power during stance phase, especially the single-support period, is lower in humans than bipedal chimpanzees, 2) the limb joint work and power required for limb swing is lower in humans than in bipedal chimpanzees, and 3) the estimated total mechanical power during walking, accounting for the storage of passive elastic strain energy in humans, is lower in humans than in bipedal chimpanzees. Humans and bipedal chimpanzees were compared at matched dimensionless and dimensional velocities. Our results indicate that humans walk with significantly less work and power output in the first double-support period and the single-support period of stance, but markedly exceed chimpanzees in the second-double support period (i.e., push-off). Humans generate less work and power in limb swing, although the species difference in limb swing power was not statistically significant. We estimated that total mechanical positive ‘muscle fiber’ work and power were 46.9% and 35.8% lower, respectively, in humans than bipedal chimpanzees at matched dimensionless speeds. This is due in part to mechanisms for the storage and release of elastic energy at the ankle and hip in humans. Further, these results indicate distinct heel strike and lateral balance mechanics in humans and bipedal chimpanzees, and suggest a greater dissipation of mechanical energy through soft tissue deformations in humans. Together, our results document important differences between human and bipedal chimpanzee walking mechanics over a full stride, permitting a more comprehensive understanding of the mechanics and energetics of chimpanzee bipedalism and the evolution of hominin walking.

## 1. Introduction

As humans, we are unique among apes and other primates in the musculoskeletal design of our lumbar column, pelvis and lower limbs. In particular, we possess a mobile lower back, a short pelvis and long lower limbs with short feet. This is distinct from the relatively immobile lumbar column, tall pelvis and short hind limbs with long feet characteristic of living African apes (e.g., Schultz, 1930, 1961; Ward, 1993; Pilbeam and Lieberman, 2017). Our limb muscle masses are more proximally located and are made up of relatively short muscle fibers and long distal limb tendons as compared to African apes (e.g., Thorpe et al., 1999; O’Neill et al., 2013, 2017). Among other capabilities, these traits are thought to facilitate humans walking with a stable pelvis on adducted, extended lower limbs (O’Neill et al., 2015), using as much as 75% less energy per distance walked than a bipedal chimpanzee (Rodman and McHenry, 1980; Sockol et al. 2007). Understanding when and where in a full stride lower/hind limb mechanics differ between human and bipedal chimpanzee walking is important for elucidating how our interspecific differences in musculoskeletal structure affect joint loading, muscle-tendon behavior and the metabolic cost of walking.

Over a walking stride, humans generate a coordinated sequence of three-dimensional (3-D) moments and power that with some exceptions, are minimal near the middle single-support period and contribute to a burst in positive power during the second double-support period (i.e., push-off), with significant moments and power continuing throughout limb swing. At the hip, in particular, there are substantial non-sagittal plane moments, work and power over a stride (e.g., Eng and Winter, 1995), comprehensive measurements of which are needed for accurate determination of skeletal loading (e.g., Stansfield et al., 2003), muscle-tendon force and fascicle length change (e.g., Arnold and Delp, 2011). Across the limb, the instantaneous 3-D joint moments, work and power reflect the net contributions from active muscle-tendon units and passive, soft tissues at each joint throughout the full gait cycle. While it is difficult to decompose the individual contributions of a mobile lower back, a short pelvis, and long lower limbs and short feet via joint-level analyses alone, these traits can be expected to act together to reduce the total mechanical work and power output of an erect human walking stride when compared to facultative bipeds, such as bipedal chimpanzees.

In addition to our skeletal structure, our more proximal limb segment masses and long, thin distal limb tendons can be expected to further reduce the total mechanical work and power output in a walking stride. It has long been hypothesized that more proximal lower limb-mass distribution reduces the moment of inertia, and therefore the mechanical power required for limb swing (e.g., Gray, 1968; Hildebrand, 1985), including in fossil hominins (Jungers and Stern, 1988). Human limb mass perturbations provide some support for this view, demonstrating that a distal shift of the mass-specific moment of inertia increases the limb swing mechanical power and metabolic costs of walking (Royer and Martin, 2005; Browning et al., 2007). Still, the magnitude of these perturbations was large, and the resulting changes in limb segment inertial properties can exceed the difference between humans and African apes or some Old World monkeys (e.g., Isler et al., 2006; Supplementary Online Material [SOM] Table S1).

Long, thin tendons, as well as other soft tissues (e.g., ligaments, aponeuroses), have a significant capacity for passive elastic strain and reducing the mechanical work and power required of muscle fibers (e.g., Alexander, 1988; Roberts, 2016). While often associated with bouncing gaits, the storage and release of elastic strain energy in human walking is now well established. Direct measurements of ankle plantar flexor muscle-tendon behavior have shown that the Achilles tendon is stretched during single-support period and then recoils in the second double-support period, making a substantial contribution to positive ankle power at push-off (e.g., Fukunaga et al., 2001; Ishikawa et al., 2005). However, elastic mechanisms in human walking are not limited to the ankle. The hip joint produces a large component of the total limb mechanical work and power output in walking (e.g., Eng and Winter, 1995; Farris and Sawicki, 2012a; Schache et al., 2015), and a portion of this is derived from non-muscular tissues stretched during the single-support period, including hip joint ligaments, associated tissues and aponeuroses (e.g., Whittington et al., 2008; Eng et al., 2015). Although there is extensive quantitative data on human walking, little comparative data exist that permit a direct assessment of whether these presumed musculoskeletal adaptations interact to reduce the total mechanical work and power output as compared to a biped that lacks this lower back, pelvis and lower limb structure, such as bipedal chimpanzees. Indeed, the absence of comprehensive, speed-controlled comparisons has left the unique aspects of human walking mechanics still not well defined, despite their critical significance for interpreting skeletal trait evolution in hominins.

This is not to suggest that data on bipedal chimpanzee walking mechanics are unknown. A seminal series of lab-based studies provided the first quantitative comparison of human and bipedal chimpanzee walking mechanics in the 1970s and 1980s (i.e., Yamazaki et al., 1979; Okada, 1985; Yamazaki, 1985). These studies include important, initial insights into the ground forces and joint moments of human and facultative bipedal walking in chimpanzees and other taxa; however, the experimental design (i.e., mismatched speeds, markerless joint kinematics) as well as the absence of variance metrics permits only limited interspecific inferences. Indeed, forces and moments are velocity-dependent parameters (e.g., Farris and Sawicki, 2012a; Schache et al., 2015; see also SOM) and, as such, interspecific comparisons are well known to be sensitive to speed. More careful, speed-controlled comparisons of human and bipedal chimpanzee walking have examined two-dimensional (2-D) sagittal plane stance phase joint moments (Sockol et al. 2007; see also Pontzer et al., 2009), but exclude joint work and power as well as limb swing. Thus, a comprehensive characterization of the 3-D moments, work and power of human and bipedal chimpanzee walking over a full stride is still unavailable.

Here, we evaluate how human and bipedal chimpanzee musculoskeletal structure affects their 3-D moments, work, and power output. Specifically, we provide the first comprehensive description and direct, speed-controlled comparisons of stride-to-stride and intraspecific variation of human and bipedal chimpanzee ground reaction forces, as well as the lower/hind limb joint moment and power profiles of each species over a full walking stride. Based on their differences in musculoskeletal structure, we then test whether: 1) the estimated limb joint work and power during stance phase, especially the single-support period, is lower in humans than bipedal chimpanzees, 2) the limb joint work and power required for limb swing is lower in humans than in bipedal chimpanzees, and 3) the estimated total mechanical power during walking, accounting for the storage of passive elastic strain energy in humans, is lower in humans than in bipedal chimpanzees. For completeness, the 3-D mechanics of bipedal chimpanzees and humans were compared at matched dimensionless (i.e., relative-speed match) and dimensional (i.e., absolute-speed match) speeds. Matched dimensionless speed comparisons minimize the effects due to differences in body size or walking speed, while emphasizing those arising specifically from differences in musculoskeletal design between humans and chimpanzees. The dimensional comparison, in contrast, permits an assessment of how sensitive the interspecific differences in 3-D mechanics are to walking speed.

## 2. Materials and methods

### 2.1. Human and chimpanzee subjects

Three-dimensional kinematic and ground force data were collected for three humans (*Homo sapiens*: age = 24.3 ± 2.3 years, mass = M_b_: 79.2 ± 6.2 kg) and three chimpanzees (*Pan troglodytes*: age = 5.5 ± 0.2 years, mass = M_b_: 26.5 ± 6.7 kg) during bipedal walking. All subjects were male. Bipedal chimpanzees walked across an 11 m rigid, level runway at self-selected speeds, following an animal trainer offering a food reward. Human data were then collected during walking along a 20 m rigid, level runway at speeds matching the chimpanzee dataset in dimensionless (i.e., relative-speed match) and dimensional (i.e., absolute-speed match) forms (Fig. 1; Table 1; SOM Table S2) using an instantaneous speed. Bipedal chimpanzees were trained for 1 hour d^-1^, 3–5 days wk^-1^, for at least six months prior to the start of data collection. The University of Massachusetts Amherst Institutional Research Board and the Stony Brook University Institutional Animal Care and Use Committee approved all human (SPHHS-HSRC-13-36) and chimpanzee (2009-1731-FAR-USDA) experiments, respectively. Prior to data collection, the human subjects each provided written informed consent.

**Figure 1.**
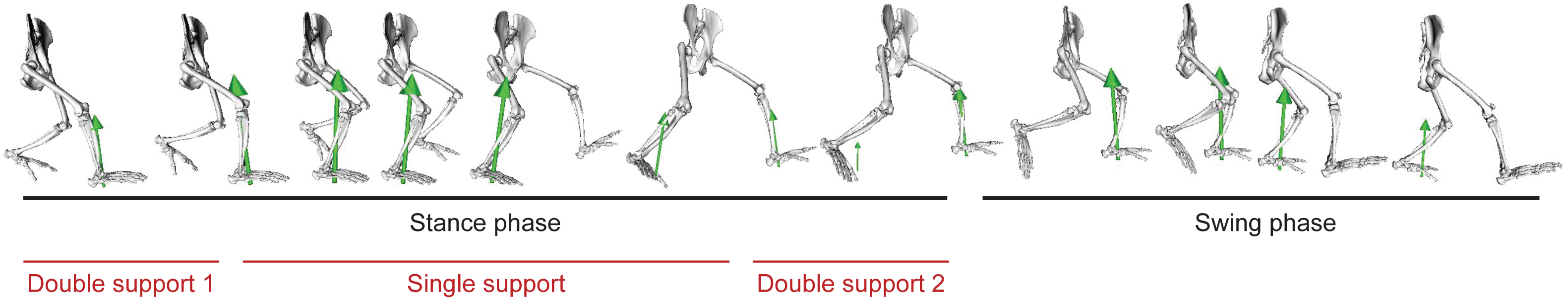
A full bipedal walking stride. A full stride includes both stance and swing phases. The stance phase is divided among the first double support (double support 1), single support, and the second double-support (double support 2) periods. In the first double-support period the right hind limb is the leading limb, while in the second double-support period the right hind limb is the trailing limb. The green arrow shows the direction and relative magnitude of the resultant ground reaction force under each foot over a full stride.

**Table 1.**
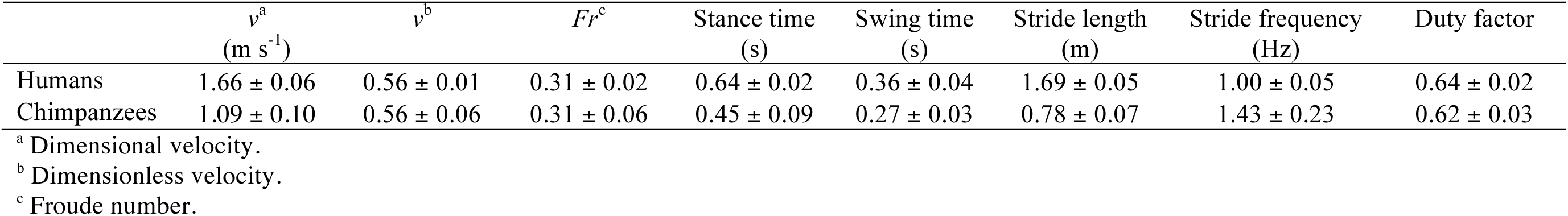
Spatiotemporal gait parameters in human and chimpanzee bipedal walking. Human data are matched to the chimpanzee data in dimensionless (i.e., relative-speed match) form (mean ± s.d.).

### 2.2. Three-dimensional marker and ground reaction force data

The procedures used for marker data collection and 3-D joint kinematic measurements in humans and chimpanzees have been detailed in O’Neill et al. (2015). In brief, marker positions were recorded using synchronized high-speed video cameras. Marker data for the humans were collected using an eleven-camera system recording at 240 Hz (Qualisys, Inc.; Gothenburg), while chimpanzee data were collected using a four-camera system recording at 150 Hz (Xcitex, Inc., Woburn). The calibrated video recording volume for the humans was created using a wand-based nonlinear transformation approach, while the recording volume for the chimpanzees was established using direct linear transformation and a custom-built calibration frame. Marker locations were digitized using Qualisys Track Manager software (Qualisys, Inc.; Gothenburg) for the humans and ProAnalyst software (Xcitex, Inc., Woburn) for the chimpanzees.

Ground reaction forces were collected using floor-mounted force platforms set into the runway flush to its surface, within the calibrated video recording volume. Two force platforms (AMTI, Watertown) sampling at 2400 Hz were used to record individual limb forces from humans, whereas an array of four force platforms (AMTI, Watertown) sampling at 1500 Hz were used to collect individual limb forces in chimpanzees throughout a bipedal walking stride. Marker data and force platform data were collected using a manual trigger that started and stopped video and force platform recording simultaneously. The *x*-, *y*-, and *z*-force data were low pass filtered with a cut-off frequency set to 60 Hz, while the marker data were filtered at 4–6 Hz. The force data were then downsampled to 240 or 150 Hz to match the synchronous video data for each species using MATLAB v. R2015 (The Mathworks, Inc., Natick). The *x*-, *y*-, and *z*- data from each force platform was transformed into the marker global coordinate system using markers positioned on the corner of each platform during the calibration of the video recording volume, prior to the start of an experiment. This permitted spatial transformations between the global coordinate system and the local force plate coordinate systems.

### 2.3. Musculoskeletal models and inverse dynamics

Three-dimensional kinematics and ground force data were processed using generic musculoskeletal models of the pelvis and lower/hind limbs of an adult human (Delp et al., 1990) and an adult chimpanzee (O’Neill et al., 2013) scaled to the size of each subject using anatomical segment markers (O’Neill et al., 2015). All models included skeletal geometry of the pelvis and lower/hind limbs relevant to determining 3-D mechanics of the hip, knee, and ankle (talocrual) joints.

An inverse kinematics algorithm was used to determine the 3-D coordinates of the scaled model of each subject of each species over the full gait cycle (O’Neill et al., 2015). This was done through a least-squares minimization of the discrepancy between the experimentally determined marker positions and the corresponding marker positions on the scaled model, subject to constraints enforced by the anatomical models of the joints (Lu and O’Connor, 1999; Delp et al., 2007). The 3-D kinematics and force data were combined with the scaled musculoskeletal model of each subject and used to calculate the net joint moments and mechanical power associated with the joint moments. Inverse dynamics combines rigid-segment linear and rotational motion, ground reaction forces, and limb segment inertial parameters to calculate net forces and moments at each joint. Detailed limb segment inertial parameters are provided for the adult human and adult chimpanzee models in the supplemental materials (SOM Table S1). These generic parameters were scaled to the size of each subject based on subject-specific measurements of body mass and segment lengths.

The instantaneous joint powers were computed at the *j*^th^ joint over both the stance and swing phase of walking as the dot product of the *j*^th^ joint moment and joint angular velocity vectors. Because the joint angles and joint moments were both expressed in the anatomically relevant but non-orthogonal joint reference frames, the moment and angular velocity vectors were both transformed into a common orthogonal reference frame before computing the dot product. This transformation is especially critical when the segments forming a joint are not closely aligned with the global reference frame, as in the abducted, flexed-limb walking of bipedal chimpanzees (O’Neill et al., 2015). The hip, knee, and ankle powers were then multiplied by two to represent the work done by both limbs over a full stride:

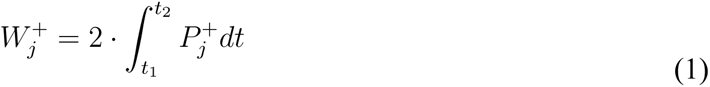

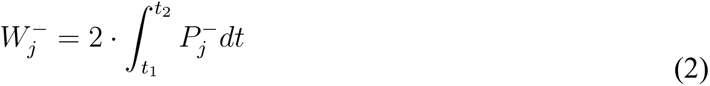

The positive and negative mechanical work of the *j*^th^ joint was computed by integrating over the time duration *t*_1_ to *t*_2_ of a walking stride, where *t*_1_ and *t*_2_ are the start and end of the first double-support period (ds1), single-support period (ss), second-double support period (ds2) of stance phase or the limb swing phase (sw). The total limb positive and negative work in the *i*^th^ period or phase of a stride were calculated as the sum of the positive and negative work at each of the *n* joints (*n* = 3 for each species):

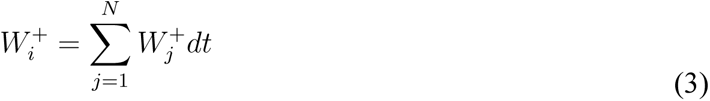

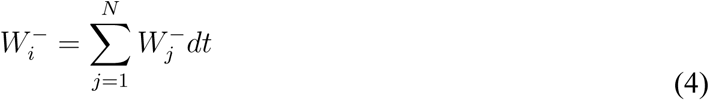

Total positive and negative mechanical work for the complete stride (*W*^+^_tot_, *W*^-^_tot_) were calculated as the sum of the positive or negative work for the first double-support period (*W*^+^_ds1_; *W*^-^_ds1_), single-support period (*W*^+^_ss_; *W*^-^_ss_), second double-support period (*W*^+^_ds2_; *W*^-^ _ds2_) and limb swing (*W*^+^_sw_; *W*^-^_sw_). The positive and negative work terms were subsequently divided by the stride time to determine average power output (*P*^+^*_i_*; *P*^-^*_i_*).

### 2.4. Elastic and ‘muscle fiber’ work and power estimates

In human walking, passive elastic energy is stored at the ankle and hip joints during the single-support period and released in the second-double-support period as well as, perhaps, in early limb swing (e.g., Fukunaga et al., 2001; Ishikawa et al., 2005; Whittington et al., 2008; Eng et al., 2015). The structures most responsible for this are the Achilles tendon at the ankle and the joint capsule ligaments and iliotibial tract at the hip. A similar mechanism is not expected at the knee joint due to the opposite phasing (i.e., positive power in single-support and negative power in the second-double-support period) of the joint power during late stance phase. We estimated that elastic energy was stored during the portion of the single-support period in which the relevant joint power curve was negative. We assumed that the elastic energy storage equaled the negative work, and that all of the elastic energy stored was supplied to the subsequent period of positive power generation. At both the dimensionless and dimensional walking speeds, the positive power generation occurred during the double-support period for the ankle, or the double-support period and early limb swing for the hip. As such, the net positive elastic work during the single-support period was estimated as:

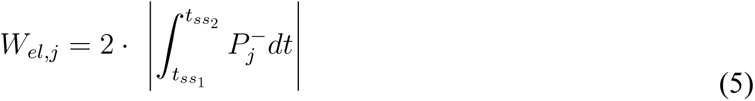

where *W*_el,*j*_ is the elastic work absorbed at the *j*^th^ joint (*a*: ankle; *h*: hip) during the single-support period of the stance phase of a walking stride, and used to generate positive elastic recoil. The positive elastic power output was computed by dividing the positive elastic work by stride time. Therefore, the total ‘muscle fiber’ work and power were the estimated positive elastic work at the ankle (*W_el,a_*) and hip (*W_el,h_*) subtracted from the total positive mechanical work (*W*^+^_tot_). The positive mechanical power attributable to ‘muscle fiber’ was computed by dividing the ‘muscle fiber’ work by stride time.

### 2.5. Analysis and statistics

Four strides per subject were analyzed for the humans (H) and chimpanzees (C). All 3-D moment and power data were normalized to 101 points over one full stride, facilitating comparisons of multiple strides. This also permitted the mean ± standard deviation (s.d.) of the moment and power curves to be determined per species.

For the chimpanzees, walking speed was calculated as the average of the instantaneous forward velocity of four markers (i.e., 3 pelvis, 1 hip marker) over the full stride, while for humans a marker placed over the sacrum was used. To account for differences in body size among subjects and species, analyses were performed with dimensionless variables, using the base units of body mass *M*_b_, gravitational acceleration *g*, and average lower/hind limb length *L*. Lower/hind limb length was measured as the average height of the greater trochanter marker from the ground during quiet standing for humans and during the middle of the single-support period for chimpanzees (H = 0.92 ± 0.05, C = 0.39 ± 0.02 m). Velocity (*v*) was made dimensionless by the divisor (*gL*)^0.5^ and as the Froude number (*Fr* = *v*^2^/*gL*). Force was made dimensionless by the divisor *M*_b_*g*, moment and work by *M*_b_g*L* and power by *M*_b_*g*^1.5^*L*^0.5^. In all cases, stance, swing, and stride duration were determined based on the ground reaction forces.

To compare the stride-to-stride, intraspecific, and interspecific variation of the ground reaction forces of our human and chimpanzee samples, the adjusted coefficient of multiple correlation (CMC) was used (Kadaba et al. 1989, r^2^_a_). The CMC represents the correlation of the ground reaction force among strides for each individual (i.e., stride-to-stride variation) or among the individuals (i.e., intraspecific variation). The balanced human and chimpanzee datasets permit a direct, interspecific comparison of CMCs computed using the species means.

To compute the difference in dimensionless work and power between humans and bipedal chimpanzees, the percent difference was calculated as |(*s*_1_ – *s*_2_)|/[(*s*_1_ + *s*_2_)/2]·100, where s_1_ (species 1) and s_2_ (species 2) are the species mean work and power parameters. Because the sample was balanced, a two-level nested analysis of variance (ANOVA) was used to compare work and power output in each period or phase of a walking stride (i.e., double-support 1, single support, double-support 2, limb swing; Fig. 1), as well as the total mechanical work and power output over a full stride to test hypotheses 1-3. A two-level nested approach allowed repeated individual subject trials to be incorporated as random effects and was therefore preferred to an independent samples t-test based on subject means. Statistical significance was set at α = 0.05. The nested ANOVA was computed in R v. 4.0.2 using the base package for Mac OS (R Development Core Team, 2020).

Due to the small sample sizes for each species, omega-squared (ω^2^) was also used to assess the effect size of the difference between chimpanzees and human comparisons outlined in hypotheses 1-3. Effect size descriptive thresholds were ω^2^ ≤ 0.01 = ‘small’, ω^2^ ≤ 0.06 = ‘medium’, ω^2^ ≤ 0.14 = ‘large’ (Field, 2013). Effect size calculations were implemented in R v. 4.0.2 using the ‘effectsize’ package (R Development Core Team, 2020).

## 3. Results

The average walking speeds of humans and chimpanzees were 1.66 ± 0.06 and 1.09 ± 0.10 m s^-1^, respectively. These correspond to similar dimensionless velocities (*v’*; H = 0.56 ± 0.01, C = 0.56 ± 0.06) and *Fr* numbers (H = 0.31±0.02, C = 0.31 ± 0.06) (Table 1). These speeds are close to—but a bit faster than—the preferred overground speeds for human walking, and well below the expected walk-run transition speed for terrestrial mammals (i.e., *v’* = 0.7; *Fr* = 0.5; Alexander, 1989; Kram et al., 1997).

Here, we describe the three-dimensional ground reaction forces, lower/hind limb joint moment and power profiles of humans and bipedal chimpanzees walking over a full stride at matched dimensionless speeds. The limb-joint work and power in stance, swing and over a full stride are determined to test hypotheses 1-3.

### 3.1. Ground reaction forces, stride-to-stride, and intraspecific variation

The ground reaction forces of humans and chimpanzees were different in both magnitude and pattern (Fig. 2). The vertical force was monophasic in bipedal chimpanzees, lacking the second peak from the single-support period into the second double-support period that is characteristic of human walking. The bipedal chimpanzee anterior-posterior force was distinct from humans, as it exhibited a smaller peak force in both posterior and anterior directions, while the mediolateral force was larger, most notably during the single-support period.

**Figure 2.**
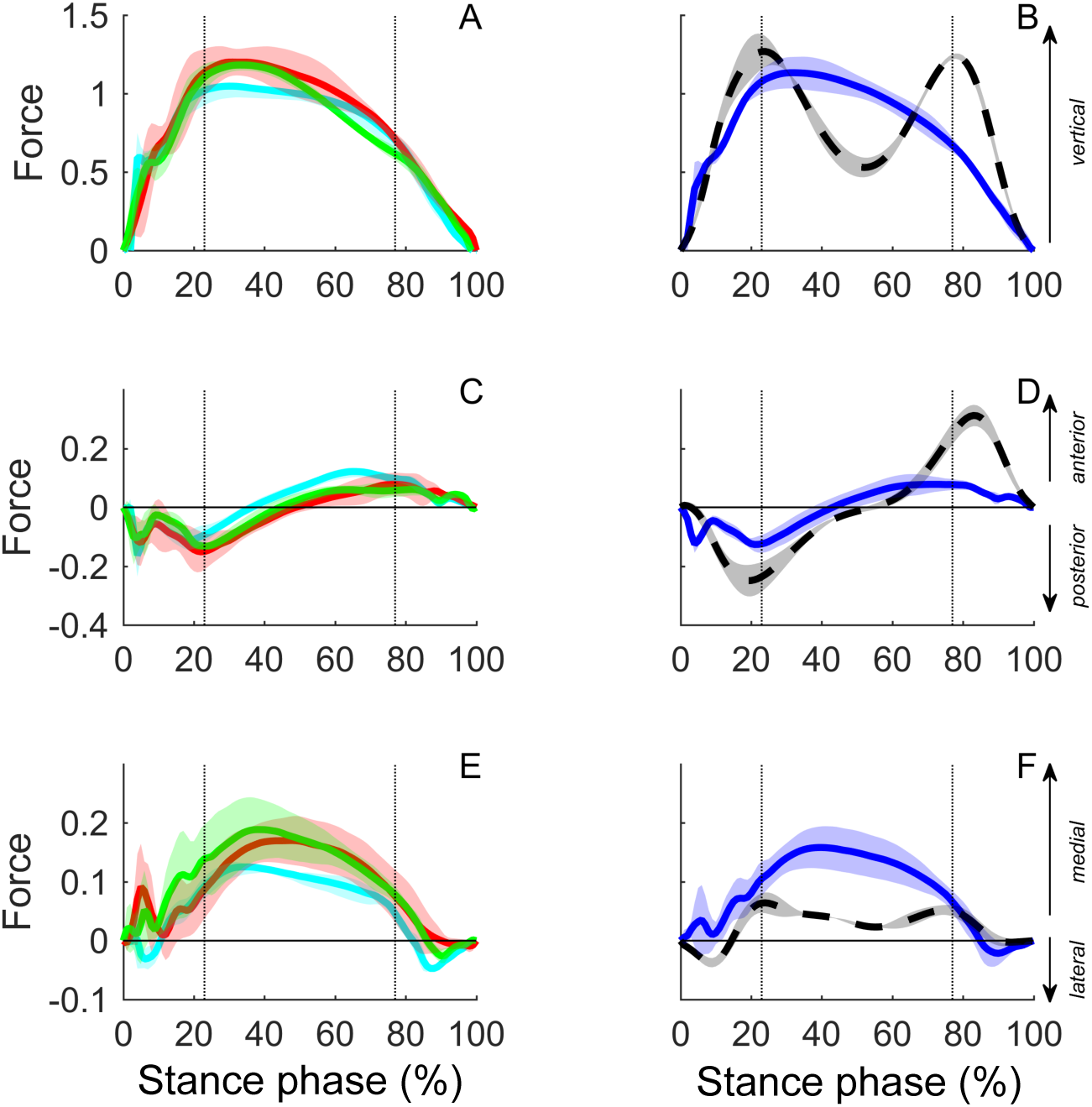
The vertical (A–B), anterior-posterior (C–D) and medio-lateral (E–F) ground reaction forces over a walking step (mean ± s.d.) for the three chimpanzees (column 1) as well as chimpanzees (solid blue line) and humans (dashed black line) as groups (column 2). Each step begins at heel strike (i.e., foot touchdown) and ends at toe off of the same foot. Broken vertical lines show the average step event times for the chimpanzees, representing contralateral limb toe off (H = 21%, C = 23%) and the contralateral limb heel strike (H = 81%, C = 77%), which define the double-support and single support periods of a step.

The CMCs were smaller between strides than between individuals for both humans and bipedal chimpanzees (Table 2). This indicates that there is more variation among humans and chimpanzees in ground reaction forces than there is within a given individual from one stride to the next. Among individuals, the medio-lateral force was the most variable in chimpanzees (r^2^_a_ = 0.847 ± 0.012) and humans (r^2^_a_ = 0.916 ± 0.038). Directly comparing the CMC values of human and chimpanzee samples indicates that, on average, chimpanzees are more variable in their ground reaction force stride-to-stride than humans at matched dimensionless speeds (H = r^2^_a_ = 0.985, C = r^2^_a_ = 0.922).

**Table 2.** Adjusted coefficient of multiple correlation (CMC) of ground reaction forces (F_y_, F_x_, F_z_).

| | $F_{y, \text{vert}}$ | $F_{x, \text{ant-post}}$ | $F_{z, \text{med-lat}}$ |
| --- | --- | --- | --- |
| Humans<sup>a</sup> | $0.994 \pm 0.003$ | $0.996 \pm 0.001$ | $0.965 \pm 0.017$ |
| Among humans<sup>b</sup> | $0.958 \pm 0.018$ | $0.972 \pm 0.007$ | $0.916 \pm 0.038$ |
| Chimpanzees<sup>a</sup> | $0.972 \pm 0.021$ | $0.911 \pm 0.054$ | $0.884 \pm 0.074$ |
| Among chimpanzees<sup>b</sup> | $0.962 \pm 0.016$ | $0.883 \pm 0.031$ | $0.847 \pm 0.012$ |

Human data are matched to the chimpanzee data in dimensionless (i.e., relative-speed match) form (mean ± s.d.). CMC mean ± s.d. for 4 trials per subject, 3 subjects. The y, x, z components of the ground reaction forces are vertical (vert), anterior-posterior (ant-post) and medial-lateral (med-lat).
<sup>a</sup> Correlations between strides within subjects.
<sup>b</sup> Correlations between subjects.

### 3.2. Limb-joint moment profiles

As expected, humans and chimpanzees were quite different in the magnitude and pattern of the limb joint moments over a full stride (Fig. 3; Table 3). At the hip, chimpanzees exhibit the largest joint moments in hip flexion-extension, with peak hip extension moment occurring in the first double-support period and continuing throughout the single-support period of stance. Unlike humans, for whom the net flexion moment begins near midstance, the net flexion moment does not occur until the second-double support period in bipedal chimpanzee walking. Humans and chimpanzees were opposite in phase in the frontal plane, where chimpanzees exhibit a net hip adduction moment throughout stance that peaks in the second-double support period. Near the middle of the single support period, the hip adduction-abduction moment approaches zero in both taxa. Chimpanzees exhibit substantial hip rotational moments, with a large internal rotation moment throughout the single-support and second double-support periods. During limb swing, chimpanzees exceed humans in the magnitude of internal and external rotational moments.

**Figure 3.**
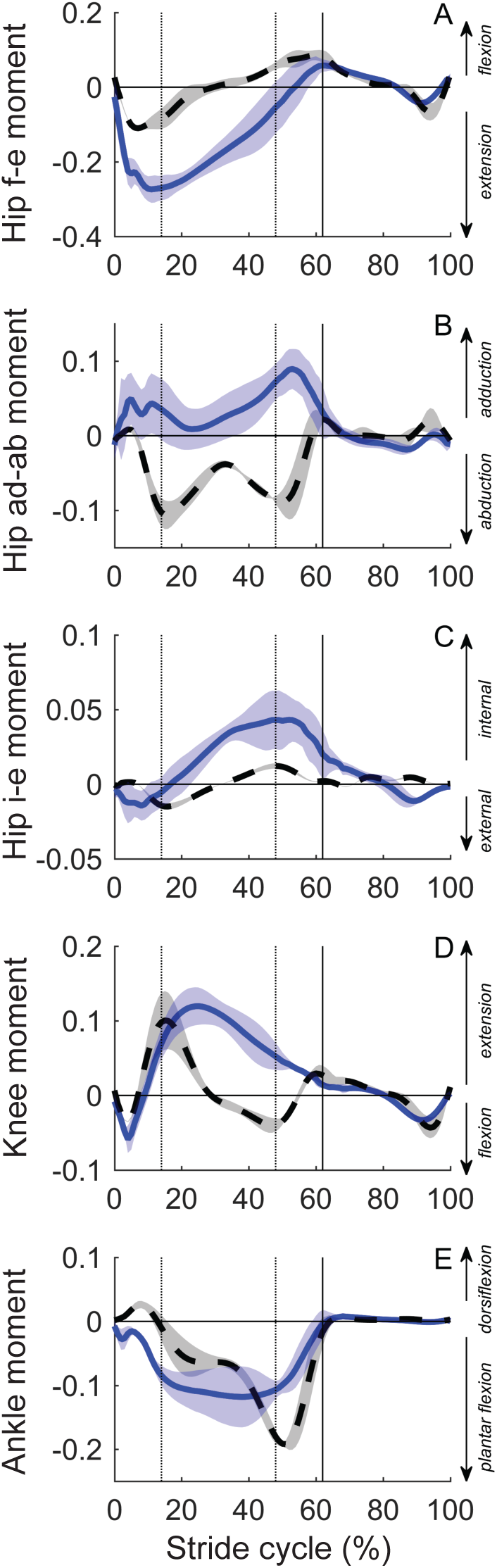
The hip flexion-extension moment (A), abduction-adduction moment (B), internal-external rotation (C), knee flexion-extension moment (D) and ankle plantar flexion-dorsiflexion moment (E) over a walking stride (mean ± s.d.) for chimpanzees (solid blue line) and humans (dashed black line). Moments are in dimensionless units. Each stride begins and ends at ipsilateral ‘heel strike’ (i.e., foot touchdown). All vertical lines show the average stride event times for the chimpanzees, which were similar among the species. The broken vertical lines represent contralateral limb toe-off (H = 13%, C = 14%) and contralateral limb ‘heel strike’ (H = 51%, C = 48%), which define the double-support and single support periods of a stride; the solid vertical line represents ipsilateral toe off (H = 63%, C = 62%), which defines the start of the swing phase.

**Table 3.**
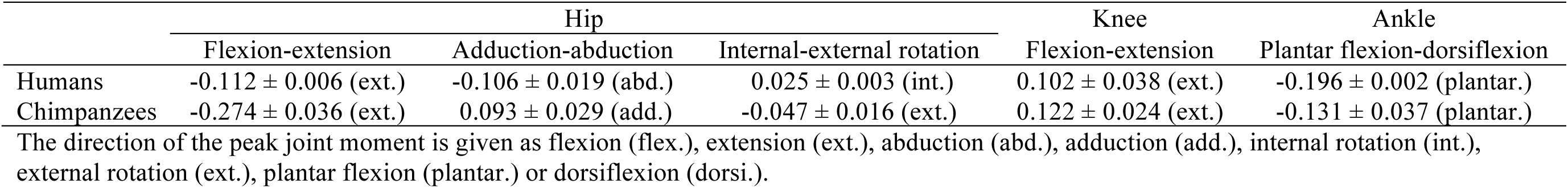
Peak joint moments in bipedal chimpanzees and humans in dimensionless units. Human data are matched to the chimpanzee data in dimensionless (i.e., relative-speed match) form (mean ± s.d.).

Humans and chimpanzees both exhibit an initial knee flexion moment in the first double-support period, and then rapidly shift to a large net extension moment at the start of the single-support period. Unlike humans, chimpanzees have a net knee extension moment throughout the single and second double-support periods. The pattern and magnitude of knee flexion-extension moments in limb swing are similar for both species.

The chimpanzee exhibits a net ankle plantar flexion moment throughout stance phase, including during the first double-support period. This differs from humans, who exhibit a net dorsiflexion moment in the first double-support period, likely associated with our distinct ‘heel strike’. Whereas chimpanzees maintain a near-constant ankle plantar flexion moment throughout the remainder of stance, there is an increase in the magnitude of the net plantar flexion moment throughout the single and second double-support periods in humans, which peaks in the second-double support period. For both species, there is a small net dorsiflexion moment throughout limb swing, on average.

### 3.3. Limb-joint power profiles

The magnitude as well as pattern of joint powers differed between humans and bipedal chimpanzees (Fig. 4; Table 4). In both humans and chimpanzees, there was negative hip power in the first double-support period. In the single-support period, there is an initial period of hip energy generation followed by hip energy absorption in humans, whereas chimpanzee hips generate energy throughout this period. In the second-double support period, there was positive power in human hips, whereas there was negative power in chimpanzee hips. For humans, the second double-support period positive power continues into limb swing, ending with a small burst of energy absorption. Chimpanzees generate substantial positive hip power in the first half of limb swing, ending with a small burst of energy generation.

**Figure 4.**
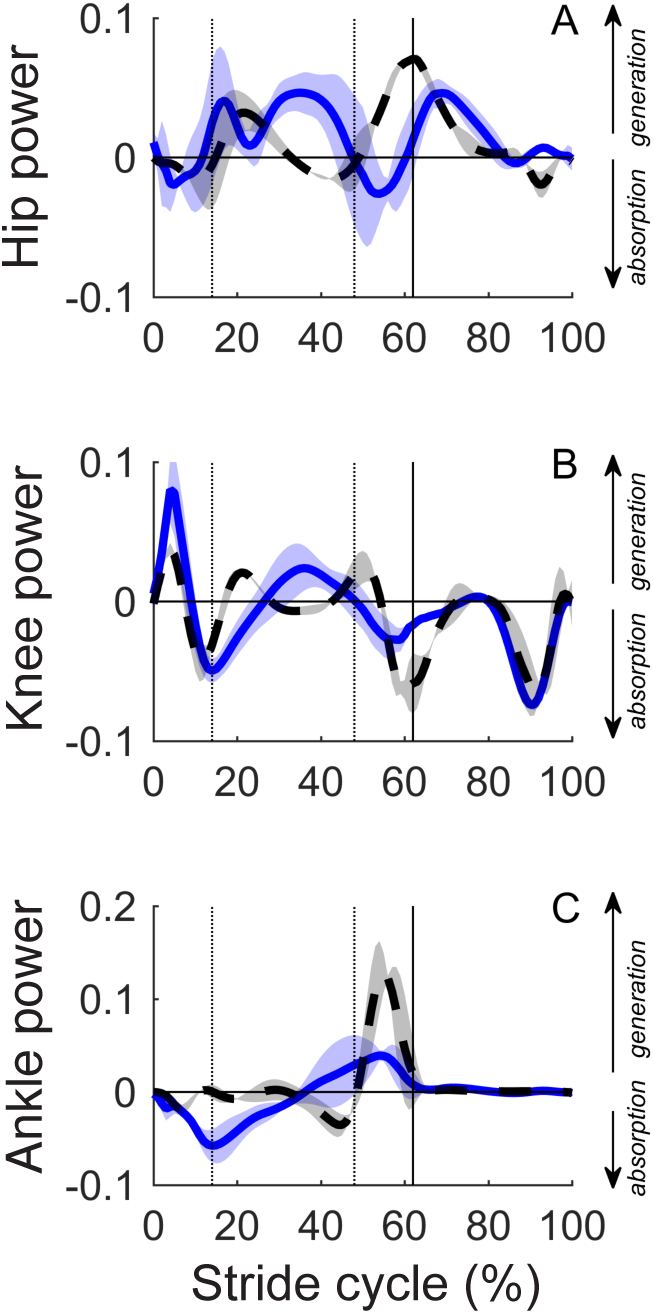
The 3-D hip power (A), knee power (B), and ankle power output (C) over a walking stride (mean ± s.d.) for humans (dashed black line) and chimpanzees (solid blue line). All vertical lines show the average stride event times for the chimpanzees, which were similar among the species. The broken vertical lines represent contralateral limb toe-off (H = 13%, C = 14%) and contralateral limb ‘heel strike’ (H = 51%, C = 48%), which define the double-support and single support periods of a stride; the solid vertical line represents ipsilateral toe off (H = 63%, C = 62%), which defines the start of the swing phase. Powers are in dimensionless units.

**Table 4.** Peak joint power in bipedal chimpanzees and humans at matched dimensionless speeds in dimensionless units. Human data are matched to the chimpanzee data in dimensionless (i.e., relative-speed match) form (mean ± s.d.).

|  | Hip power |  | Knee power |  | Ankle power |  |
| --- | --- | --- | --- | --- | --- | --- |
|  | + | − | + | − | + | − |
| Humans | $0.071 \pm 0.010$ | $-0.031 \pm 0.002$ | $0.038 \pm 0.004$ | $-0.078 \pm 0.005$ | $0.145 \pm 0.037$ | $-0.034 \pm 0.011$ |
| Chimpanzees | $0.067 \pm 0.016$ | $-0.050 \pm 0.016$ | $0.083 \pm 0.027$ | $-0.076 \pm 0.004$ | $0.052 \pm 0.014$ | $-0.059 \pm 0.018$ |

At the knee joint, there was a substantial positive power peak in the first double-support period in both chimpanzees and humans, followed by complex, cyclical pattern of energy absorption and generation through the single-support and second double-support periods. In the middle of the single-support period, chimpanzees have a small burst of energy generation, whereas humans are near zero power output. Throughout limb swing, there was substantial negative power at the knee in both species, especially in the second half of swing phase.

The ankle joint in chimpanzees is characterized by energy absorption followed by energy generation over the stance phase. This is distinct from humans, where a small amount of energy absorption occurs in the first double-support period, followed by a period of negative power at the end of the single-support period and then substantial positive power output in the second-double support period. The peak dimensionless positive power in the second-double support period is almost three times greater in humans than chimpanzees at matched dimensionless walking speeds, on average (Table 4). For both species, there is a relatively small amount of energy generation from the ankle throughout limb swing.

### 3.4. Limb-joint work and power in stance (hypothesis 1)

Humans generate less dimensionless positive mechanical work (*W*^+^_ds1_: F_1,18_ = 21.38, *p* = 0.0099; ω^2^ = 0.77) and power (*P*^+^_ds1_: F_1,18_ = 12.67, *p* = 0.0236; ω^2^ = 0.66) than chimpanzees during the first double-support. The same was true for the single-support period positive mechanical work (*W*^+^_ss_: F_1,18_ = 27.58, *p* = 0.0063; ω^2^ = 0.82) and power (*P*^+^_ss_: F_1,18_ = 13.51, *p* = 0.0213; ω^2^ = 0.68). In the second double-support period, humans generate much more positive mechanical work (*W*^+^_ds2_: F_1,18_ = 38.15, *p* = 0.0035; ω^2^ = 0.86) and power than chimpanzees (*P*^+^_ds2_: F_1,18_ = 76.51, *p* = 0.0009 ω^2^ = 0.93; Fig. 5). In all cases, the effect sizes were ‘large’. This provides partial support for our first hypothesis.

**Figure 5.**
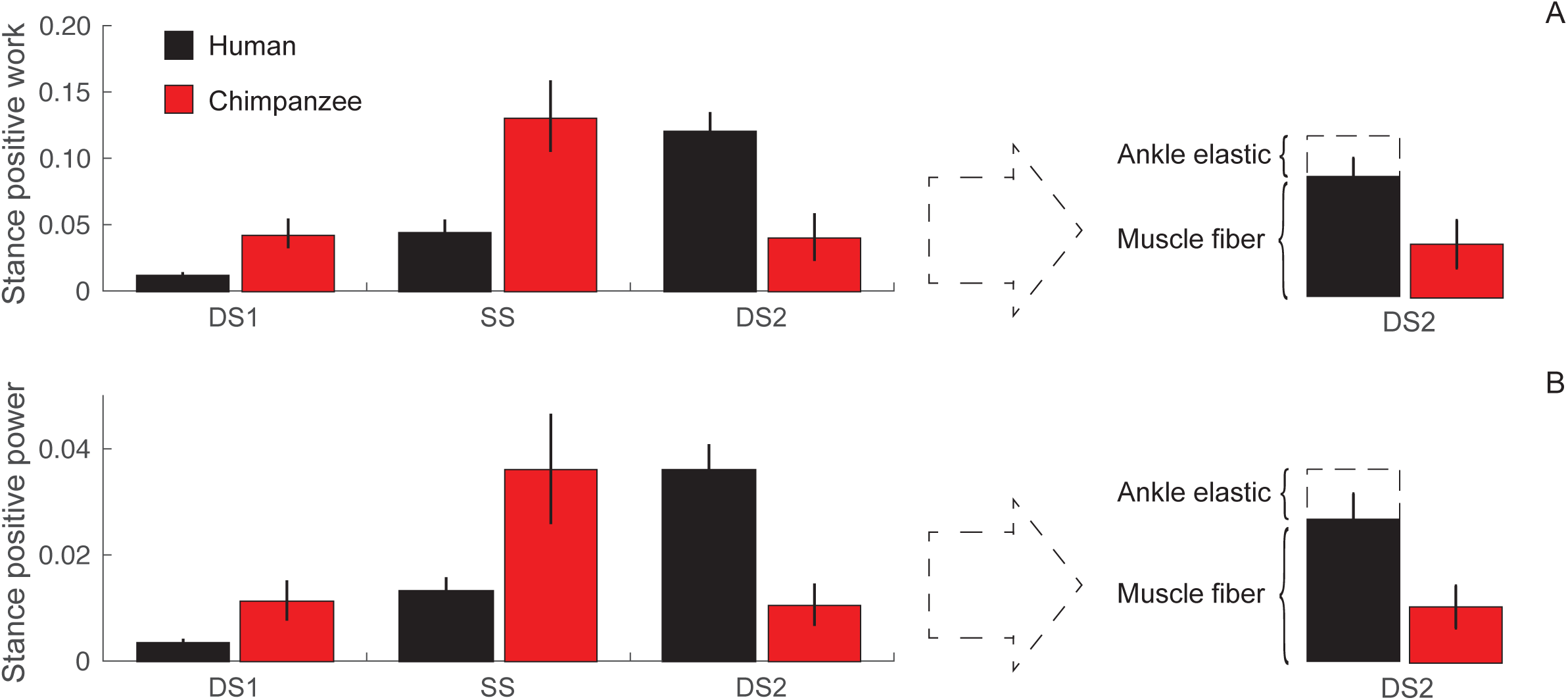
The stance phase positive joint work (A) and power (B) output in the first double support (DS1), single support (SS) and second double-support (DS2) period of a walking stride for humans (black) and chimpanzees (red) (mean ± s.d.). The relative contributions of the ankle elastic work and power (‘ankle elastic’) and muscle fiber work and power (‘muscle fiber’) to the total limb work and power output in the human DS2 period is shown.

### 3.5. Limb-joint work and power in swing (hypothesis 2)

Humans generate less dimensionless positive mechanical work (*W*^+^_sw_: F_1,18_ = 16.71, *p* = 0.0016; ω^2^ = 0.72) and power (*P*^+^_sw_: F_1,18_ = 6.649, *p* = 0.0614; ω^2^ = 0.48) than chimpanzees in limb swing. The difference in positive work was significant, but the difference in power only near significant; however, in both cases the effect size was ‘large’ (Fig. 6). This provides partial support for our second hypothesis.

**Figure 6.**
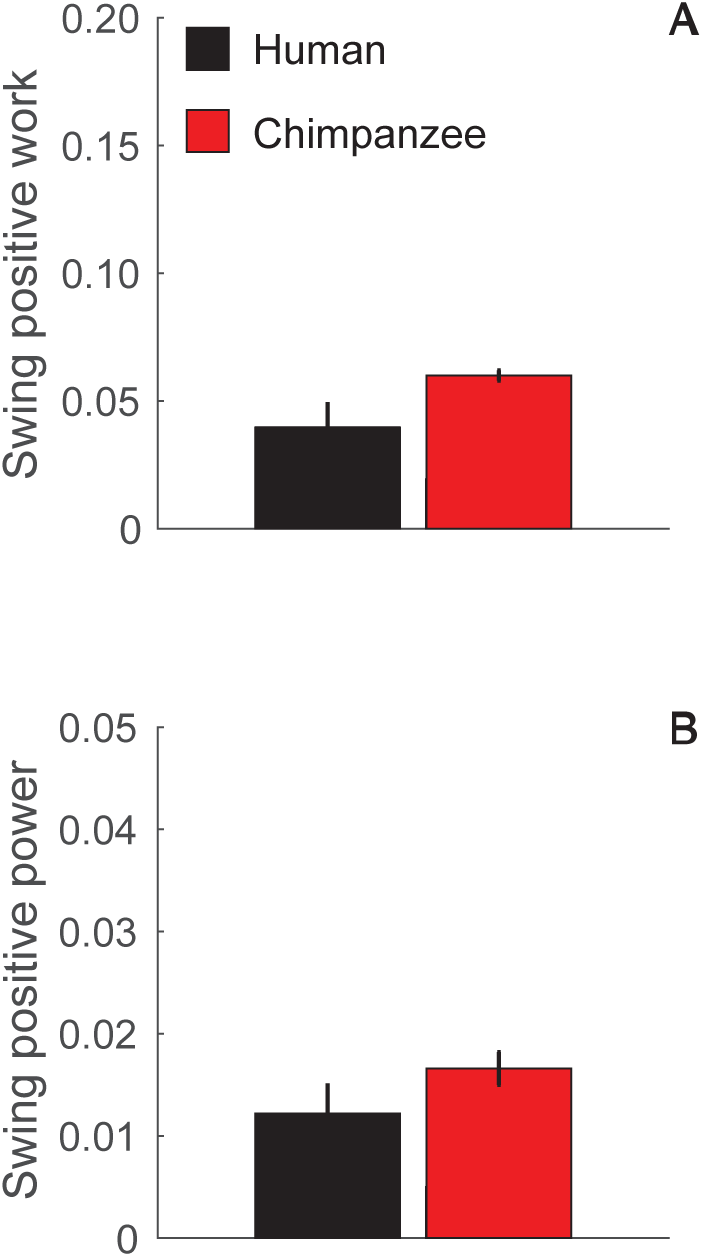
Total positive limb work (A) and average positive limb power (B) used to swing the limb during walking in humans (black) and chimpanzees (red; mean ± s.d.).

### 3.6. Total mechanical work and power (hypothesis 3)

The total dimensionless positive mechanical work (*W*^+^_tot_) and power (*P*^+^_tot_) was 23.3% and 12.0% smaller in humans than chimpanzees, respectively. The difference in mechanical work was ‘large’ and significant (F_1,18_ = 11.55, *p* = 0.0273; ω^2^ = 0.64), while the difference in power was a ‘medium’ effect, and not statistically significant (F_1,18_ = 1.372, *p* = 0.3070; ω^2^ = 0.06). The modest differential in total mechanical power is due to the substantial positive power output in the second double-support period of a human walking stride. However, when the mechanical work and power attributable to elastic storage and release at the human ankle and hip were accounted for, the estimated total positive ‘muscle fiber’ work and power percent differential between species increased to 46.9% and 35.8%, respectively. The differences in estimated total positive ‘muscle fiber’ work (F_1,18_ = 33.19, *p* = 0.0045, ω^2^ = 0.84) and power (F_1,18_ = 10.56, *p* = 0.0314, ω^2^ = 0.61) between species were both significant, and the effect sizes were ‘large’ (Fig. 7), supporting our third hypothesis.

**Figure 7.**
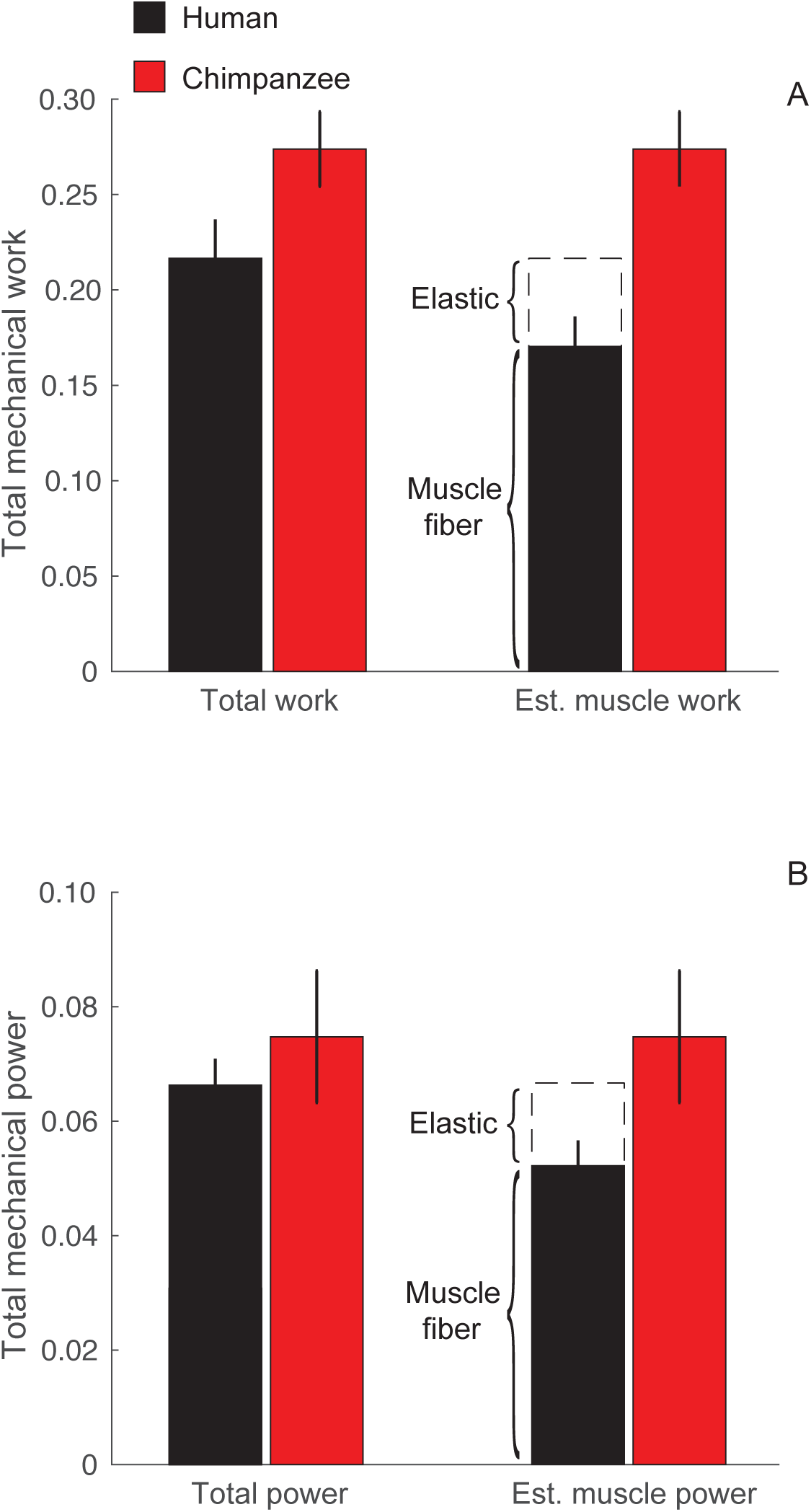
Total positive limb work (A) and average positive limb power (B) over a full walking stride in humans (black) and chimpanzees (red; mean ± s.d.). The human total lower limb work and power is due to contributions from both elastic mechanisms at the hip and ankle (‘elastic’) as well as muscle fibers (‘muscle fiber’).

### 3.7. Distribution of work across limb joints

In human walking, the stance phase work is predominately from the ankle (47.0 ± 6.5%), and from the hip in swing phase (82.9 ± 3.2%). This contrasts with chimpanzees in which most of their positive mechanical work is from the hip in both stance (49.1 ± 3.7%) and swing (88.5 ± 1.9%) phases (Fig. 8).

**Figure 8.**
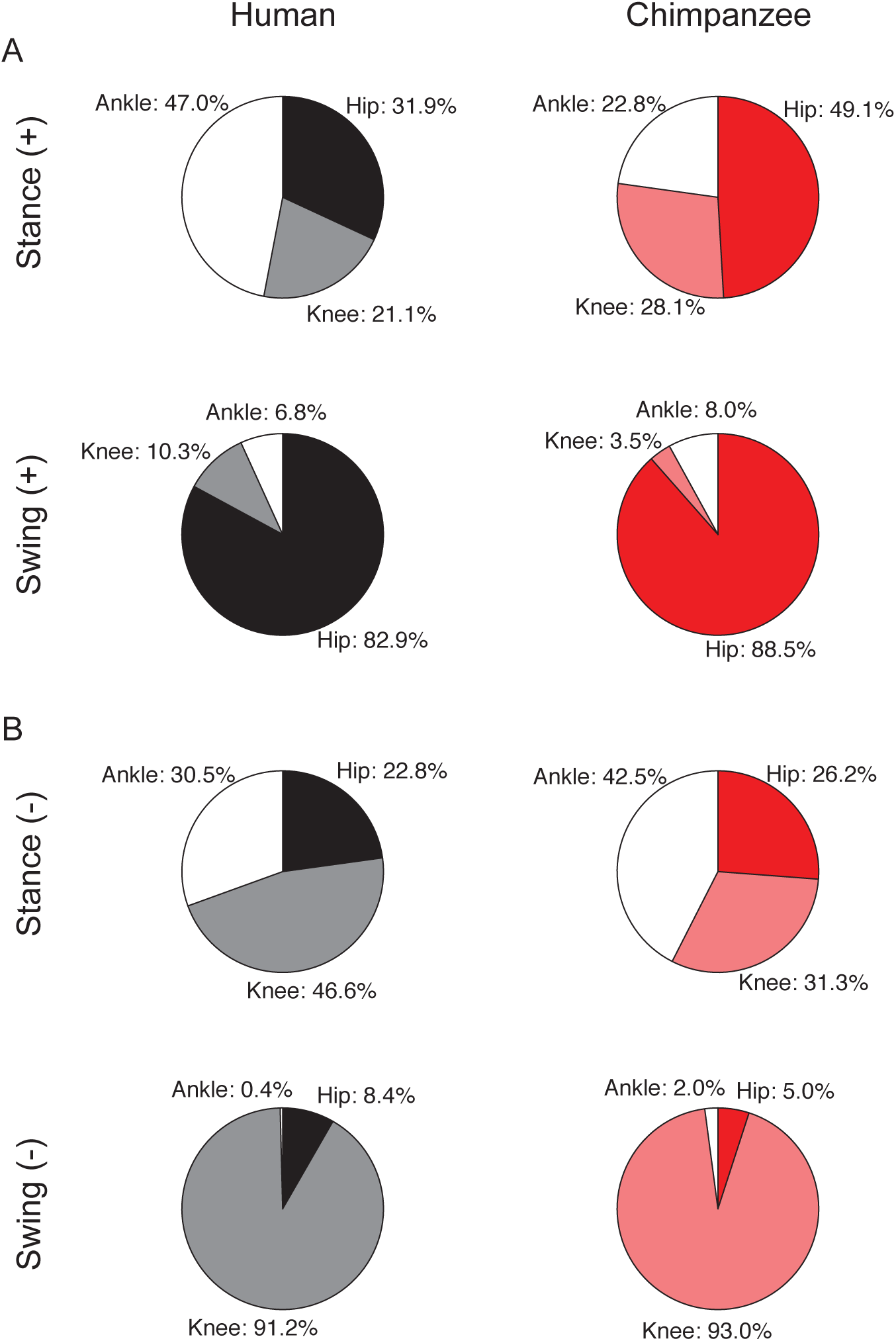
The distribution of positive mechanical work (A) and negative mechanical work among the limb joints (B) during stance and swing in humans (column 1) and chimpanzees (column 2).

In stance phase, human negative work is predominately absorbed at the knee (46.6 ± 3.0%), whereas chimpanzees absorb most of their mechanical work at the ankle (42.4 ± 2.0%). In swing phase, the knee accounts for nearly all of the negative work in both species (H = 91.2 ± 4.6%; C = 93.0 ± 2.1%).

### 3.8. Dimensional vs. dimensionless walking mechanics

For the comparison at matched dimensional speeds (i.e., absolute-speed match), the average human walking speed was 1.08 ± 0.02 m s^-1^. This corresponds to a *v’* of 0.36 ± 0.01 and *Fr* of 0.13 ± 0.02 (SOM Table S2), which are slower than the relative-speed match.

Overall, the humans had smaller moments, work and power at matched dimensional speed than at matched dimensionless speed. As such, the interspecific differences in dimensional positive mechanical work and power output were consistently larger, with the exception of the second double-support period positive work and power (SOM Figs. S2–S8; SOM Tables S2–S5). In this case, the positive mechanical work and power output were more similar between species than at the relative-speed match (SOM Fig S5). The human 3-D joint moments and powers had similar profiles at 1.08 ms^-1^ and 1.66 ms^-1^. However, the magnitude of the interspecific difference in total mechanical work and power were much larger (H = 83.6% vs. C = 77.8%; SOM Fig. S7) than at matched dimensionless speeds, even without accounting for elastic power output. Indeed, the interspecific difference in total ‘muscle fiber’ work and power are larger still (H = 106.9% vs. C = 111.9%; SOM Fig S7). This reinforces the importance of careful, speed-controlled comparisons for testing the effects of musculoskeletal structure on locomotor capabilities.

## 4. Discussion

### 4.1. Ground reaction forces, stride-to-stride and intraspecific variation

Our results are broadly consistent with previous descriptions of the 3-D ground force components in bipedal chimpanzee walking (Kimura et al., 1977; Pontzer et al., 2009). As expected, all three animals exhibited monophasic vertical force patterns, with small magnitude anterior-posterior force peaks compared to a human walking step at similar dimensionless speeds, especially in the second double-support period (i.e., push-off). The species difference in peak vertical force magnitude (H = 1.27, C = 1.13; Fig. 2) supports the inference that facultative bipeds walk with lower peak vertical force (Schmitt, 2003), albeit by a small amount at matched dimensionless speeds. However, this difference is not present at matched dimensional speeds (SOM Fig. S2). Overall, the ground reaction forces in bipedal chimpanzees appear quite similar to the 3-D forces of ‘highly-trained’ bipedal macaques (Kimura et al., 1977; Ogihara et al., 2007), as well as some other facultative bipeds (Kimura, 1977). A qualitative comparison of our chimpanzee dataset to similar data from ‘highly-trained’ bipedal macaques (Ogihara et al., 2007) suggests that bipedal chimpanzees and macaques are much more similar to each other than either are to humans.

The data herein indicate that bipedal chimpanzees exceed humans in the variation of their ground reaction forces from stride to stride. Among individuals, chimpanzees are more variable than humans in their anterior-posterior and medio-lateral ground forces, but similar in their vertical component. These results are consistent with CMC results for the pelvis and lower/hind limb 3-D kinematics reported in O’Neill et al. (2015). The similarities in the vertical force CMC values suggest that the monophasic pattern is quite consistent across our three subjects, as suggested by average patterns from chimpanzees in earlier studies from a range of ages and body masses (Kimura et al., 1977; Pontzer et al., 2009).

### 4.2. Joint moments

Our results are consistent with earlier 2-D data on the sagittal plane moments of humans and bipedal chimpanzees throughout stance (Yamazaki et al., 1981; Sockol et al., 2007). In particular, bipedal chimpanzees—like all facultative bipeds studied to date (Yamazaki et al., 1981; Yamazaki, 1985)—have a substantial hip extension moment for nearly the full duration of stance.

However, these 3-D data also provide insight into bipedal chimpanzee mechanics in the non-sagittal planes, and over a full walking stride. Hip abduction-adduction moments are different in magnitude and opposite in direction for human and bipedal chimpanzee walking. The absence of an abductor moment in the stance phase in bipedal chimpanzee walking, and throughout the single-support period in particular, provides quantitative support to the longstanding view that the human ‘lateral balance mechanism’ is a derived aspect of hominin walking (e.g., Stern and Susman, 1981; Lovejoy, 1988). The marked contrast in hip abduction-adduction moments between humans and bipedal chimpanzees demonstrates that the single-support period of a bipedal walking stride need not follow the simple ‘mechanical model’ of hip loading, which assumes a large abduction moment during single-support (e.g., Lovejoy et al., 1973; Ruff, 1995; see also Warrener, 2017). This is an important consideration in general, but especially when extending this human-based approach to the earliest hominins (e.g., Richmond and Jungers, 2008).

The single-support period of bipedal chimpanzees is characterized by a larger internal (i.e., medial) rotation moment than in human walking. This supports the inference of Stern and Susman (1981:164) that for chimpanzees, “… the problem posed during the single support phase of bipedality is to prevent collapse into lateral rotation at the stance side hip.” Our results confirm and extend this inference, indicating that there is also a substantial hip adduction moment in bipedal chimpanzee walking, which reaches a similar magnitude to the internal rotation moment near the end of the single-support period. Taken together, the moment data herein confirm that the distinction between the 3-D mechanics of the single-support period in bipedal chimpanzee and human walking is a reliance on ‘rotation-based’ or ‘rotation-adduction-based’ hip mechanics in chimpanzees, and ‘abduction-based’ hip mechanics in humans for ‘maintaining lateral balance’. How individual hind-limb muscle-tendon units contribute to balancing these moments in bipedal chimpanzees requires additional quantitative analyses, since an individual muscle-tendon unit will, in general, have non-zero moment arms about all three hip joint axes that can change in magnitude and sign during movement (e.g., O’Neill et al., 2013).

The joint moment data also extend our earlier findings (O’Neill et al., 2015, 2018) that the use of a ‘heel strike’ during walking in humans is distinct, not just in its consistency (Elftman and Manter, 1938; Webber and Raichlen, 2016), but in both its associated ankle kinematics and mechanics, despite some qualitative similarities with other African apes overall (Schmitt and Larson, 1993). At the human ankle, there is a consistent, net dorsiflexion moment from ‘heel strike’ through the first double-support period, which contrasts with the net plantar flexor moment in bipedal chimpanzees. How these different foot strike patterns impact ankle muscle-tendon mechanics and energetics in walking is still unclear, but some limited metabolic data indicate that evolution of the hominin ‘heel strike’ is of much greater consequence for walking than running (e.g., Cunningham et al., 2010; Gruber et al., 2013). When human-like ‘heel strike’ walking first appeared in hominin evolution is still unclear, although a robust calcaneal tuber and lateral plantar process has been used to infer its presence as early as *Australopithecus afarensis* (e.g., Latimer and Lovejoy, 1989; Prang, 2015; DeSilva et al., 2019). Still, this is difficult to establish even from fossil footprints such as the Laetoli site tracks (e.g., Leakey and Hay, 1979; Masao et al., 2016), since both human and bipedal chimpanzee foot strikes induce relatively deep heel impressions (Hatala et al., 2016a).

Human and bipedal chimpanzee joint moments are broadly similar in direction and magnitude throughout limb swing, except in hip internal-external rotation. This is consistent with greater circumduction of the hind limb in bipedal chimpanzee walking (O’Neill et al., 2015). The limb-swing rotational moments also provide some indirect support to the inference of Stern and Larson (1993:425) that m. obturator externus is activated “to counteract [a] medial rotatory effect” in the first half of limb swing, as there is a net internal (i.e., medial) rotation moment at this time in bipedal chimpanzees.

### 4.3. Mechanical work and power in stance

Our results provide some support for the first hypothesis that human walking involves lower work and power output in stance phase than bipedal chimpanzees. This is due, in large part, to significantly less positive mechanical work and power output in the first double-support and single-support periods of a human walking stride. While both humans and chimpanzees absorb mechanical energy at the hip in the first double-support period, there is greater net positive mechanical work at the knee (Figs. 3A, B) in bipedal chimpanzee walking.

The greater positive mechanical work in the single-support period in chimpanzees is due primarily to greater positive power at the hip, with small contributions from the knee and ankle. This is consistent with previous assessments of sagittal plane hip moments, which had indicated a substantial force-requirement at the hip during stance phase in bipedal chimpanzees (Sockol et al., 2007; Pontzer et al., 2009). In middle single-support, the lower back, pelvis and lower limbs of humans facilitate near-zero lower limb joint power, followed by a short period in which mechanical energy is absorbed at the hip and ankle (i.e., ‘preload’: Kuo et al., 2005; Zelik et al., 2015).

### 4.4. Elastic work and power

In contrast to the first double-support and single-support periods, humans markedly exceed bipedal chimpanzees in total positive mechanical work and power during the second double-support period due to substantial mechanical energy generation at the ankle for push-off. Muscle-tendon models (e.g., Hof et al., 1983) and direct imaging of muscle fiber dynamics (e.g., Fukanaga et al., 2001; Ishikawa et al., 2005; Farris and Sawicki, 2012b) indicate that much of the positive power for human push-off is due to the elastic recoil of the Achilles tendon, stretched during the single-support period. Based on the mechanical energy absorbed at the ankle during the single-support period at this speed, we estimated that 39 ± 13% of the positive power for ankle push-off is from elastic storage, on average (Fig 5). The upper error margin of our indirect, inverse dynamics-based approximation overlaps direct imaging of muscle fiber behavior in humans walking between 1.25 and 1.75 ms^-1^, which indicates a significant contribution from elastic power (i.e., 50 to 55%), at least for m. gastrocnemius (Farris and Sawicki, 2012b). It has been reported that the typical procedures used in direct imaging studies can overestimate the elastic energy returned from tendons (Zelik and Franz, 2017), which may account for some of the difference in mean values. Of course, both these indirect and direct estimates are subject to the limitations of the inverse-dynamics approach and would benefit from continued refinement using detailed musculoskeletal models that account for individual muscle-tendon unit behavior (e.g., Hornet and Zelik, 2016).

The contribution of elastic positive hip work to the second double-support period and limb swing is less well understood but has been estimated to be 10–58% across a range of walking speeds (Yoon and Mansour, 1982; Varhas et al., 1990; Whittington et al. 2008). Herein, we estimated that the human hip stored and released 25.5 ± 10.7% of the positive work to the second double-support period and limb swing, which is in the middle of this published range. Hip joint capsular ligaments and associated tissues have long been identified as a source of hip elastic work (e.g., Neptune et al., 2008; Whittingon et al., 2008), since the anterior tissues (e.g., iliofemoral ligament) will be strained during hip extension. The iliotibial tract, which has a hip flexion moment arm, appears to contribute as well (Eng et al., 2015). Direct imaging of hip flexor muscle fiber behavior (e.g., rectus femoris) would help elucidate the role of muscle and tendon to positive power output in the second double-support and limb swing periods of a human walking stride.

Over a full stride, our estimated total elastic work contributions (i.e., ankle and hip: 24.2 ± 3.1% at 1.66 ms^-1^; 31.1 ± 7.2% at 1.08 ms^-1^) is similar to the series elastic element contributions to the total lower limb work from individual muscle tendon dynamics (e.g., 29.4% tendon elastic work to muscle fiber work at 1.2 ms^-1^: Sasaki et al. 2009). This suggests that our estimates are reasonable initial approximations. It is well known that net joint-level measures do not equate to total muscle-tendon unit work and power, owing to bi-articular muscle recruitment and muscle co-contraction (Sasaki et al. 2009), which may differ between human and bipedal chimpanzee walking.

The hind limb joint mechanics and muscle-tendon structure of bipedal chimpanzees suggest a limited capacity for elastic work and power output in a walking stride. First, the joint mechanics data herein demonstrate that, unlike human walking, the second half of the single-support period of chimpanzee bipedal walking involves no net negative hip or ankle work (see Fig. 4 and SOM Fig. S4) for elastic energy storage and is instead a continuous period of positive work. The absence of this negative work suggests that there is no significant elastic contribution from the hip and ankle to ‘push-off’ in bipedal chimpanzees. This is also implied by the hip flexion-extension range of motion, which is about half that of humans (O’Neill et al., 2015: C: 27°; H: 48°), indicating less capacity for hip tissue strain and elastic energy storage overall. Indeed, the passive tissue strain and elastic work of the iliotibial tract of bipedal chimpanzees has been estimated to be only a small fraction of human values (Eng et al., 2015), due in large part to this kinematic difference. Second, chimpanzee pelvis and hind limb muscles are also characterized by relatively longer muscle fibers than human lower limb muscles, and the near absence of a free Achilles tendon (e.g., Thorpe et al., 1999; O’Neill et al., 2013, 2017). Chimpanzee pelvis and hind limb relative muscle fiber lengths (i.e., ratio of muscle fiber length to total muscle-tendon unit length) are about 15% longer than humans overall (C = 0.59 ± 0.21, H = 0.44 ± 0.25), and about twice as long for the leg/shank muscles in particular (C = 0.33 ± 0.14, H = 0.15 ± 0.04; Arnold et al., 2010; O’Neill et al., 2013; see also Thorpe et al., 1999). This architectural design requires the muscle fibers to more closely track the muscle-tendon length changes due to the joint motions, implying a limited capacity for tendon strain and thus storage and release of elastic energy. Still, short, stiff tendons, long, thin internal tendons and sarcomeric proteins may permit some degree of elastic work and power output (e.g., Alexander, 1988; Roberts, 2016). Decomposition of joint-level 3-D mechanics among individual muscles and series elastic elements via optimization can provide a more detailed accounting of the muscle-tendon mechanical work and power (e.g., Anderson and Pandy, 1993, 2001), and more precise estimates of the elastic contribution to human and bipedal chimpanzee walking.

### 4.5. Mechanical work and power in limb swing

Our results also provide some limited support for our second hypothesis that human limb structure reduces limb swing positive work and power. The greater positive work and, to a lesser extent, power in bipedal chimpanzee limb swing is consistent with the hypothesized differences in limb structure. However, it is difficult to isolate the contribution of the limb-mass distribution per se. This is because bipedal chimpanzees engage in more hip rotation about abducted, flexed hind limbs throughout limb swing (O’Neill et al. 2015), for reasons likely independent of limb-mass distribution. Increasing limb flexion (Wu and Kuo, 2016) and circumduction (Shorter et al., 2017) in limb swing increases limb joint work and positive power over a stride in human perturbation studies, regardless of limb-mass distribution.

Elastic work and power from the hip likely contribute to both the second double-support period and the first half of limb swing in human walking. Given the sinusoidal shape of the positive power burst at the hip that peaks at the stance-swing transition (Fig. 5A), we can estimate that about half of the hip elastic work and power is output for limb swing, which would further increase the human-chimpanzee limb swing differential. In contrast, in the second double-support period of bipedal chimpanzee walking, the hip is insufficiently extended (i.e., 25° of hip flexion, on average; O’Neill et al., 2015) to strain the associated elastic tissues and permit a positive elastic power contribution to the first half of limb swing. This suggests a distinct contribution of human hip mechanics during the single and second double-support periods to reducing limb swing work and power, independent of limb-mass distribution.

### 4.6. Total mechanical work and power

On average, human walking involves 23.3% and 12.0% less positive mechanical work and power than bipedal chimpanzees, respectively. These differentials are enhanced by human capacities for elastic work and power output over a stride, which are assumed to be zero in bipedal chimpanzees herein. Together, these results support our third hypothesis, as the total ‘muscle fiber’ work and power output of the human lower limb is 46.9% and 35.8% lower than the hind limbs of bipedal chimpanzees walking at matched dimensionless speeds. This is consistent with the broad-sense expectation that our mobile lower back, pelvis and long lower limbs with short feet of humans reduce the total mechanical work and power output of our walking stride when compared to bipedal chimpanzees; however, decomposing the individual contributions of specific traits to these joint-level differences is complex, and requires additional investigation.

Our total mechanical power differential accounts for some, but not all, of the differential in the metabolic cost of walking between humans and bipedal chimpanzees (i.e., 50 to 75%: Sockol et al., 2007; Pontzer et al., 2014). Of course, joint-level metrics do not account for differences in muscle co-contraction, muscle contractile behavior (i.e., force-length-velocity dynamics), muscle architecture and myosin heavy chain content between species (e.g., O’Neill et al., 2013, 2017), all of which can further increase the human-chimpanzee differential. For example, the relatively small, mobile limb joints of chimpanzees (e.g., Jungers, 1988) may require higher levels of muscle co-contraction for joint stabilization over a bipedal walking stride. In our view, model-based approaches that permit optimization of individual muscle mechanics and metabolic costs can improve upon previous joint-level ‘force-based approximations’ of muscle energy use (e.g., Roberts et al., 1998; Sockol et al., 2007; Pontzer et al., 2009), since they parameterize individual-muscle excitation patterns as well as the metabolic costs of activation, cross-bridge cycling, shortening-lengthening and associated contractile element dynamics (e.g., Umberger and Rubenson, 2011).

### 4.7. Distribution of work and power across the limb joints

In humans, positive mechanical work and power output has been shifted to the distal joints, with 47% of the work performed at the ankle throughout stance phase. Notably, this facilitates the use of shorter muscle fibers (Thorpe et al., 1999; Arnold et al., 2010; O’Neill et al., 2013, 2017) and a greater reliance on elastic energy storage and release at the ankle. Both humans and chimpanzees have relative muscle fiber lengths (i.e., ratio of muscle fiber length to total muscle-tendon unit length) that are more than twice as long for pelvis and thigh muscles (H = 0.55 ± 0.15, C = 0.67 ± 0.20) as for leg muscles (H = 0.15 ± 0.04, C = 0.33 ± 0.14; Arnold et al., 2010; O’Neill et al., 2013; see also Thorpe et al., 1999). Our enlarged m. soleus and long Achilles tendon (e.g., O’Neill et al., 2013), in particular, are likely derived traits within the hominin lineage that enhance our positive push-off mechanics as well as elastic energy storage and release. Bipedal chimpanzees have a more proximal distribution of limb work, with 49% performed at the hip during stance. A reliance on proximal joints for positive work and power necessitates the activation of longer, parallel-fibered muscles (e.g., Thorpe et al., 1999; O’Neill et al., 2013), whose shortening-lengthening behavior is expected to more faithfully track 3-D joint motion, with less capacity for decoupling of muscle fiber and tendon strain. In limb swing, both humans and bipedal chimpanzees primarily use the hip. To the extent that the longer, heavier feet of chimpanzees affect their limb swing mechanics, this may account for some of the relatively larger contribution of hip (H = 83%, C = 89%) work (Fig. 8) in bipedal chimpanzees than humans.

Over a steady, level walking stride, no net mechanical work is performed on the environment, and the total whole-body positive and negative work must sum to zero. In humans, muscle-tendon units of the lower limb provide most of the positive work through rotations of the joints; however, soft tissues such as the heel pad, intervertebral discs or articular cartilages likely compress and perform significant negative work on the body without rotating the lower limb joints (Zelik and Kuo, 2013; Zelik et al., 2015). Our results indicate that humans exhibit an imbalance of limb joint work, with average negative-to-positive ratios of 67 ± 9% per step (i.e., excluding limb swing). This is nearly identical to ratio of 68% per step in level walking in DeVita et al. (2007). At matched dimensionless speeds, bipedal chimpanzee negative-to-positive work is more balanced, with a ratio of 88 ± 8% per step. This is due to greater negative work—and more active muscle-tendon unit lengthening—in bipedal chimpanzee than human walking. Although the metabolic cost of performing negative muscle fiber work is low (Woledge et al., 1985), energy dissipation through soft tissue deformations has no cost at all. As an example, a human-like heel strike permits mechanical energy dissipation *via* the heel pad (e.g., Baines et al., 2018; Honert and Zelik, 2019), rather than muscle-tendon lengthening. The shift from facultative to habitual bipedal walking in hominin evolution may have increased our reliance on soft (non-muscular) tissues for mechanical energy dissipation, which may have some metabolic and fatigue-resistance benefits in walking overall.

### 4.8. Limitations of this study

Studies of the development of human walking mechanics indicate that children over the age of 5 years have adult-like gait mechanics and metabolic costs, once differences in size are accounted for (Sutherland, 1997; Stansfield et al., 2003; Weyand et al., 2010). Chimpanzees grow into adults at a faster rate than humans (Zihlman et al., 2004), but are still sub-adult between the ages of 5 and 6 years old. Ontogenetic studies of chimpanzee bipedal walking ground reaction forces indicated that by the age of 5, chimpanzees exhibit adult-like mechanics, independent of speed (Kimura, 1996). Sagittal plane hip, knee and ankle joint moments of bipedal chimpanzees that range in age from 6 to 33 years old were all quite similar to each other (Pontzer et al., 2014). A qualitative comparison of our ground reaction forces and sagittal plane moments during stance phase to these data indicates that our results are similar to chimpanzees across a broad age and size range and, given the commonalities in the between-subject CMCs for both the force and kinematic results (Table 2; O’Neill et al., 2015) as well, should therefore be representative of bipedal chimpanzee walking in general.

Our analyses did not quantify intrinsic foot motion. Bipedal chimpanzees walk with less 3-D midfoot (Holowka et al. 2017) and forefoot (Fernandez et al., 2016) motion than humans. These new data indicating greater intrinsic foot motion in humans than bipedal chimpanzees were unexpected given long-standing dichotomies of human feet as ‘stiff’ and chimpanzees as ‘mobile’ in bipedal walking (e.g., Elftman and Manter, 1935; Susman, 1983). However, the ‘stiff’ and ‘mobile’ dichotomy may be more reflective of passive ranges of motion as well as an overemphasis on the ‘midtarsal break’ in the single-support period of bipedal chimpanzee walking. Indeed, most of human 3-D midfoot and forefoot motion actually occurs in the second double-support period, not in the single-support period (e.g., Lundgren et al. 2003; Holowka et al. 2015). Our inverse kinematics weighted markers located on the calcaneus more heavily than other foot markers for calculating ankle joint kinematics, which keeps intrinsic foot deformation from contaminating ankle joint angular velocities, and therefore ankle joint power. This approach helps prevent overestimation of ‘push-off’ power, which is important for proper calculation of plantar flexor muscle function (e.g., Zelik and Hornet, 2018). Still, we did not take into account the possible intrinsic foot power generation in late stance. Greater human intrinsic foot motion may have some adaptive value in contributing positive power to push-off (e.g., Takahashi et al., 2017; Farris et al., 2019; but also see Zelik et al., 2015). The limited midfoot and forefoot motion, together with small peak forces in the second double-support period of bipedal chimpanzee walking (see Fig. 2) suggest little foot positive power in push-off, by comparison. Thus, the inclusion of intrinsic foot mechanics in comparisons of human and bipedal chimpanzee walking may affect the total limb power output and is an important area of future study for understanding hominin foot evolution.

### 4.9. Implications for fossil hominin bipedal biomechanics

The single-support period of a walking stride has particular relevance for hominin evolution since a number of skeletal traits have been tied to ‘maintaining lateral balance’ in hominins, such as ilia orientation (e.g., Stern and Susman, 1981; Lovejoy, 1988). Humans are well known to use an ‘abductor-based hip mechanism’ in the single-support period, an assumption that has been extended to a wide range of fossil hominins (e.g., Lovejoy et al., 1973; Ruff, 1995), including the earliest hominins (e.g., Richmond and Jungers, 2008). Our results demonstrate that bipedal chimpanzees use an alternate mechanism, in which there are large internal rotation and adduction moments in the single-support period. This ‘rotation-based’ or ‘rotation-adduction-based’ mechanism for ‘maintaining lateral balance’ confirms and extends the inferences of Stern and Susman (1981), who proposed these single-support period mechanics for bipedal chimpanzees and other apes (i.e., gibbons, orangutans). This represents an alternative means of maintaining whole-body stability during bipedal walking; one that likely preceded the ‘abductor-based’ mechanism in the hominin lineage. Indeed, the likelihood that the kinematics and mechanics of bipedal walking at the origins of hominin lineage were quite distinct from humans and much more similar to bipedal walking in facultative bipeds (cf. O’Neill et al., 2018) is an important consideration when searching for skeletal adaptations for ‘terrestrial bipedal travel’ in the earliest putative hominins (e.g., Macchiarelli et al., 2020). More experimental and modeling-simulation studies are needed to evaluate how musculoskeletal trait evolution in the pelvis and hind/lower limbs, including shifts in ilia length and orientation (e.g., Stern and Susman, 1981; Lovejoy, 1988) facilitate the use of an ‘adduction-based mechanism’ for ‘maintaining lateral balance’.

An important difference in musculoskeletal structure between humans and chimpanzees is that chimpanzees possess a longer ischium than humans (e.g., Stern and Susman, 1983; Lovejoy, 1988). A long ischium is evident in *Ar. ramidus* at 4.4. million years ago (Lovejoy et al. 2009), as well as all Miocene apes with associated pelvis remains (Hammond et al., 2013, 2020; Morgan et al., 2015; Ward, 1993; Ward et al., 2019). Komza et al. (2018:4136) has recently proposed that shortening the ischium in hominin evolution facilitated ‘hamstrings-powered hip extension’. However, our results demonstrate that the second-double support period in a human walking stride is associated with a net flexion moment at the hip, not a net extension moment (cf. Fig. 3 and SOM Fig. S3). As such, the net mechanical energy generated during the second-double support period at the human hip is not due to ‘hamstrings power’, but instead to substantial hip flexor muscle (e.g., m. iliopsoas; Capellini et al., 2006) and elastic power. Indeed, in both human and bipedal chimpanzee walking, the hamstring muscles (i.e., mm. semimembranosus, semitendinosus and biceps femoris, long head) are quiescent in late stance (e.g., Winter and Yak, 1987; Capellini et al., 2006; Schmitz et al., 2009; Larson and Stern, unpublished data). Instead, our results indicate that a shorter ischium in humans is associated with greater hip extension than bipedal chimpanzees in the second half of the single-support and double-support periods, which permits a passive elastic contribution at the hip, thereby contributing positive power through push-off into limb swing.

Our results demonstrate significant differences between humans and bipedal chimpanzees over a full walking stride but place new emphasis on the second half of the single support period (i.e., for elastic tissues strain) and the second-double support period (i.e., for push-off) for the evolution of hominin walking. An abducted hallux in *Ar. ramidus* (Lovejoy et al., 2009b; Simpson et al., 2019; see also Haile-Selassie et al., 2012) suggests that the earliest hominins still utilized a relatively weak push-off in the double-support period at 4.4 Mya, quite distinct from human walking. Lovejoy et al. (2009b) has proposed that, in the absence of an adducted hallux, the center of pressure was directed along the full length of the foot and through the more lateral phalanges in *Ar. ramidus*. In bipedal chimpanzees, the center of pressure remains proximal to the phalanges over a walking step, including during the second double-support period (see Fig 1; see also Vereecke et al. 2003). Skeletal correlates of an enhanced capacity for metatarsophalangeal flexion in digits 2–4 in *Ar. ramidus* compared to African apes (Lovejoy et al., 2009b; Fernández et al., 2018; Simpson et al., 2019), raise the possibility of a more distal progression of the center of pressure than in bipedal chimpanzees; however, the lateral digits still lack the structural integrity of a large hallux for applying a substantial push-off impulse via the ankle plantar flexors (and, to a lesser extent, the intrinsic foot tissues: Takahashi et al., 2017). Nevertheless, this may represent some improvement over the limited capabilities of bipedal chimpanzees or a *Pan*-*Homo* last common ancestor with an African ape-like foot (e.g., Pilbeam and Lieberman, 2017; Prang, 2019). A more adducted hallux is evident in *Au. afarensis* (e.g., Susman et al., 1984; DeSilva et al., 2019) as well as the associated Lateoli site G and site S footprints (Leakey and Hay, 1979; Masao et al., 2016), although the substantial positive power at the ankle via a human-like push-off may not appear until *Homo* (e.g., Fernández et al., 2016, 2018). Indeed, the differences between the ca. 3.7 Mya Laetoli footprints and the 1.5 Mya Ileret footprints may reflect this (Bennett et al., 2009; Hatala et al., 2016b; but see Raichlen and Gordon, 2017). More experimental-modeling studies of the effects of foot musculoskeletal structure on push-off mechanics in walking are needed to clarify these issues.

## 5. Conclusions

The analyses presented herein provide the first description of the 3-D hip, knee and ankle moments, work, and power of bipedal chimpanzees, and direct comparisons to humans walking at matched dimensionless and dimensional speeds over a full walking stride. They provide partial or full support for our three main hypotheses relating musculoskeletal structure to mechanics across a full walking stride.

Still, it is difficult to decompose the individual contributions of musculoskeletal traits such as a mobile lower back, a short pelvis, and long lower limbs and short feet via joint-level analyses alone. Additional contributions from differences in upper body size and shape or neural control may also be relevant. Yet, this is a necessary step for linking the hominin fossil record to shifts in hominin walking capabilities. As such, future research should use the results herein to investigate the effects of isolated traits, both skeletal (e.g., ilia orientation, hallux adduction) and soft tissue (e.g., m. soleus, Achilles tendon), on the mechanics of human walking (e.g., single support, ‘push-off’, etc.) and across lower limb joints over a stride. While studies of isolated traits in hominin evolution have historically been approached using either comparative morphology or, to a much lesser extent, experimentation in living taxa (e.g., Ross et al., 2002), there is an increasingly viable and important role for computer simulation and optimization methods to establish direct cause-and-effect relationships based on high-fidelity models of the neuromusculoskeletal systems (e.g., Falisse et al., 2019). This approach may be especially useful for testing the adaptive nature of trait combinations not present in living species or along a single lineage, such as hominins. High-quality, controlled experimental data, such as the force, moment, work, and power measurements of humans and bipedal chimpanzees herein, provide the experimental basis for these studies.

## References

Alexander, R.M., 1988 Elastic Mechanisms in Animal Movement. Cambridge University Press. Cambridge, UK.

Alexander, R.M., 1989 Optimization and gaits in the locomotion of vertebrates. Physiol. Rev. 69, 1199–1227.

Anderson, F.C., Pandy, M.G., 1993. Storage and utilization of elastic strain energy during jumping. J. Biomech. 26, 1413–1427.

Anderson, F.C., Pandy, M.G., 2001. Static and dynamic optimization solutions for gait are practically equivalent. J. Biomech. 34, 153–161.

Arnold, E.M., Delp, S.L. 2011., Fibre operating lengths of human lower limb muscles during walking. Philos. Trans. R. Soc. Lond. B. Biol. Sci. 366, 1530–1539.

Arnold, E.M., Ward, S.R., Lieber, R.L., Delp, S.L., 2010. A model of the lower limb for analysis of human movement. Ann. Biomed. Eng. 38, 269–279.

Arnold, E.M., Hamner, S.R., Seth, A., Millard, M., Delp, S.L., 2013. How muscle fiber lengths and velocities affect muscle force generation as humans walk and run at different speeds. J. Exp. Biol. 216, 2150–2160.

Baines, P.M., Schwab, A.L., van Soest, A.J., 2018. Experimental estimation of energy absorption during heel strike in human barefoot walking. PLoS One 13, e0197428.

Bennett, M., Harris, J., Richmond, B., Brain, D., Mbua, E., Kiura, P., Olago, D., Kibunija, M., Omuombo, C., Behrensmeyer, A.K., Huddart, D., Gonzalez, S., 2009. Early hominin foot morphology based on 1.5-million-year-old footprints from Ileret, Kenya. Science 323, 1197–1201.

Browning, R.C., Modica, J.R., Kram, R., Goswami, A., 2007. The effects of adding mass to the legs on the energetics and biomechanics of walking. Med Sci. Sports Exerc. 39, 515–525.

Cappellini, G., Ivanenko, Y.P., Poppele, R.E., Lacquaniti, F., 2006. Motor patterns in human walking and running. J. Neurophysiol. 95, 3426–3437.

Cunningham, C.B., Schilling, N., Anders, C., Carrier, D.R., 2010. The influence of foot posture on the cost of transport in humans. J. Exp. Biol. 213, 790–797.

Delp, S.L., Loan, J.P., Hoy, M.G., Zajac, F.E., Topp, E.L., Rosen, J.M., 1990 An interactive graphics-based model of the lower extremity to study orthopaedic surgical procedures. IEEE Trans. Biomed. Eng. 37, 757–767.

Delp, S.L., Anderson, F.C., Arnold, A.S., Loan, P., Habib, A., John, C.T., Guendelman, E., Thelen, D.G., 2007 OpenSIM: Open-source software to create and analyze dynamic simulations of movement. IEEE Trans. Biomed. Eng. 54, 1940–1950.

DeSilva, J., McNutt, E., Benoit, J., Zipfel, B., 2019. One small step: A review of Plio-Pleistocene hominin foot evolution Yearbk. Phys. Anthropol. 168, 63–140.

DeVita, P., Helseth, J., Hortobagyi, T., 2007. Muscles do more positive than negative work in human locomotion. J. Exp. Biol. 210, 3361–3373.

Elftman, H., Manter, J., 1935 Chimpanzee and human feet in bipedal walking. Am. J. Phys. Anthropol. 1, 69–79.

Eng, C.M., Arnold, A.S., Biewener, A.A., Lieberman, D.E., 2015. The human iliotibial band is specialized for elastic energy storage compared with the chimp fascia lata. J. Exp. Biol. 218, 2382–2393.

Eng, J.J., Winter, D.A., 1995. Kinetic analysis of the lower limbs during walking: What information can be gained from a three-dimensional model? J. Biomech. 28, 753–758.

Falisse, A., Serrancoli, G., Dembia, C.L., Gillis, J., Jonkers, I., De. Groote, F., 2019. Rapid predictive simulations with complex musculoskeletal models suggest that diverse healthy and pathological human gaits can emerge from similar control strategies. J. R. Soc. Interface 16, 20190402.

Farris, D.J., Sawicki, G.S., 2012a. The mechanics and energetics of human walking and running: A joint level perspective. J. R. Soc. Interface 9, 110–118.

Farris, D.J., Sawicki, G.S., 2012b. Human medial gastrocnemius force-velocity behavior shifts with locomotion speed and gait. Proc. Natl. Acad. Sci. USA 109, 977–982.

Farris, D.J., Kelly, L.A., Crewsell, M.A., Litchwark, G., 2019. The functional importance of human foot muscles for bipedal locomotion. Proc. Natl. Acad. Sci. USA 116, 1645–1650.

Field, A., 2013. Discovering Statistics Using IBM SPSS Statistics, 4th edition. Sage, London.

Fernández, P.J., Holowka, N.B., Demes, B., Jungers, W.L., 2016. Form and function of the human and chimpanzee forefoot: Implications for early hominin bipedalism. Sci. Rep. 6, 30532.

Fernández, P.J., Mongle, C.S., Leakey, L., Proctor, D.J., Orr, C.M., Patel, B.A., Almécija, S., Tocheri, M.W., Jungers, W.L., 2018. Evolution and function of the hominin forefoot. Proc. Natl. Acad. Sci. USA 115, 8746–8751.

Fukunaga, T., Kubo, K., Kawakami, Y., Fukashiro, S., Kanehisa, H., Maganaris, C.N., 2001. In vivo behavior of human muscle tendon during walking. Proc. R. Soc. Lond. B Biol. Sci. 268, 229–233.

Gray, J., 1968. Animal Locomotion. W.W. Norton & Co, Inc., New York.

Gruber, A.H., Umberger, B.R., Braun, B., Hamill, J., 2013. Economy and rate of carbohydrate oxidation during running with rearfoot and forefoot strike patterns. J Appl. Physiol. 115, 194–201.

Hammond, A.S., Alba, D.M., Almécija, S., Moyà-Solà, S., 2013. Middle Miocene *Pierolapithecus* provides a first glimpse into early hominid pelvic morphology. J. Hum. Evol. 64, 658–666.

Hammond, A.S., Rook, L., Anaya, A.D., Cioppi, E., Costeur, L., Moyà-Solà, S., Almécija, S., 2020. Insights into the lower torso in late Miocene hominoid *Oreopithecus bambolii*. Proc. Natl. Acad. Sci. USA 117, 278–284.

Hatala, K.G., Demes, B., Richmond, B.G., 2016a. Laetoli footprints reveal bipedal gait biomechanics different from those of modern humans and chimpanzees. Proc. R. Soc. Lond. B Biol. Sci. 283, 20160235.

Hatala, K.G., Roach, N.T., Ostrofsky, K.R., Wunderlich, R.E., Dingwall, H.L., Villmoare, B.A., Green, D.J., Harris, J.W.K., Braun, D.R., Richomond, B.G., 2016b. Footprints reveal direct evidence of group behavior and locomotion in *Homo erectus*. Sci. Rep. 6, 28766.

Haile-Selassie, Y., Saylor, B.Z., Deino, A., Levin, N.E., Alene, M., Latimer, B., 2012. A new hominin foot from Ethiopia shows multiple Pliocene bipedal adaptations. Nature 483, 565–569.

Hildebrand M. 1985. Walking and running. In: Hildebrand, M., Bramble, D.M., Liem, K.F., Wake, D.B. (Eds.), Functional Vertebrate Morphology. Harvard University Press, Boston, pp. 38–57

Hof, A.L., Geelen, B.A., van den Berg, J.W., 1983. Calf muscle moment, work and efficiency in level walking: Role of series elasticity. J. Biomech. 16, 523–537.

Holowka, N.B., O’Neill, M.C., Thompson, N.E., Demes, B., 2017. Chimpanzee and human midfoot motion during bipedal walking and the evolution of the longitudinal arch of the foot. J. Hum. Evol. 104, 23–31.

Honert, E.C., Zelik, K.E., 2016. Inferring muscle-tendon unit power from ankle joint power during the push-off phase of human walking: Insights from a multiarticular EMG-driven model. PLoS One 11, e0163169.

Honert, E.C., Zelik, K.E., 2019. Foot and shoe responsible for the majority of soft tissue work in early stance of walking. Hum. Mov. Sci. 64, 191–202.

Ishikawa, M., Komi, P.V., Grey, M.J., Lepola, V., Bruggemann, G-P., 2005. Muscle-tendon interaction and elastic energy usage in human walking. J. Appl. Physiol. 99, 603–608.

Isler, K., Payne, R.C., Gunther, M.M., Thorpe, S.K.S., Savage, R., Crompton, R.H., 2006. Inertial properties of hominoid limb segments. J. Anat. 209, 201–218.

Jungers, W.L., 1988. Relative joint size and hominoid locomotor adaptations with implications of the evolution of hominid bipedalism. J. Hum. Evol. 17, 247–265.

Jungers, W.L., Stern, J.T., Jr., 1988. Body proportions, skeletal allometry and locomotion in the Hadar hominids: A reply to Wolpoff. J. Hum. Evol. 12, 673–684.

Kadaba, M.P., Ramakrishnan, H.K., Wooten, M.E., Gainey, J., Gorton, G., Cochran, G.V.B., 1989. Repeatability of kinematic, kinetic, and electromyographic data in normal adult gait. J. Ortho. Res. 7, 849–860.

Kimura, T., 1996. Centre of gravity of the body during the ontogeny of chimpanzee bipedal walking. Folia Primatol. 66, 126–136.

Kimura, T., Okada, M., Ishida, H., 1977. Dynamics of primate bipedal walking as viewed from the force of foot. Primates 18, 137–147.

Komza, E.E., Webb, N.M., Harcourt-Smith, W.E.H., Raichlen, D.A., D’Août, K., Brown, M.H., Finestone, E.M., Ross, S.R., Aerts, P., Pontzer, H., 2018. Hip extensor mechanics and the evolution of walking and climbing capabilities in humans, apes and fossil hominins. Proc. Natl. Acad. Sci. USA 115, 4134–4139.

Kram, R., Domingo, A., Ferris, D.P., 1997 Effect of reduced gravity on the preferred walk-run transition speed. J. Exp. Biol. 200, 821–826.

Kuo, A.D., Donelan, J.M., Ruina, A., 2005. Energetic consequences of walking like an inverted pendulum: Step-to-step transitions. Exerc. Sport Sci. Rev. 33, 88–97.

Latimer, B., Lovejoy, C.O., 1989. The calcaneus of *Australopithecus afarensis* and its implications for the evolution of bipedality. Am. J. Phys. Anthropol. 78, 369–386.

Leakey, M.D., Hay, R.L., 1979. Pliocene footprints in the Laetoli beds and Laetoli, northern Tanzania. Nature 278, 317–323.

Lovejoy, C.O., 1988 Evolution of human walking. Sci. Am. 259, 118–125.

Lovejoy, C.O., Suwa, G., Spurlock, L., Asfaw, B., White, T.D., 2009a. The pelvis and femur of *Ardipithecus ramidus*: The emergence of upright walking. Science 326, 71e1–71e6.

Lovejoy, C.O., Latimer, B., Suwa, G., Asfaw, B., White, T.D., 2009b Combining prehension and propulsion: The foot of *Ardipithecus ramidus*. Science 326, 72e1–72e8.

Lovejoy, C.O., Latimer, B.M., Spurlock, L., Haile-Selassie, Y., 2016. The pelvic girdle and limb bones of KSD-VP-1/1. In: Haile-Selassie, Y., Su, D.F. (Eds.), The Postcranial Anatomy of Australopithecus afarensis: New Insights from KSD-VP-1/1. Springer, New York, pp. 155–178.

Lu, T.W., O’Connor, J.J., 1999. Bone position estimation from skin marker coordinates using global optimization with joint constraints. J. Biomech. 32, 129–134.

Lundgren, P., Nester, C., Liu, A., Arndt, A., Jones, R., Stacoff, A., Wolf, P., Lundberg, A., 2008. Invasive in vivo measurement of rear-, mid- and forefoot motion during walking. Gait Posture 28, 93–100.

Macchiarelli, R., Bergeret-Medina, A., Marchi, D., Wood, B., 2020. Nature and relationships of *Sahelanthropus tchadensis*. J. Hum. Evol. 149, 102898.

Masao, F.T., Ichumbaki, E.B., Cherin, M., Barili, A., Boschian, G., Iurino, D.A., Menconero, S., Moggi-Cecchi, J., Manzi, G., 2016. New footprints from the Laetoli (Tanzania) provide evidence for marked body size variation in early hominins. eLife 5, e19568.

Morgan, M.E., Lewton, K.L., Kelley, J., Otárola-Castillo, E., Barry, J.C., Flynn, L.J., Pilbeam, D., 2015. A partial hominoid innominate from the Miocene of Pakistan: Description and preliminary analyses. Proc. Natl. Acad. Sci. USA 112, 82–87.

Neptune, R.R., Sasaki, K., Kautz, S.A., 2008. The effect of walking speed on muscle function and mechanical energetics. Gait Posture 28, 135–143.

Ogihara, N., Hirasaki, E., Kumakura, H., Nakatuskasa, M., 2007. Ground-reaction-force profiles of bipedal walking in bipedally trained Japanese monkeys. J. Hum. Evol. 53, 302–308.

Ogihara, N., Makishima, H., Nakatsukasa, M., 2010. Three-dimensional musculoskeletal kinematics during bipedal locomotion in the Japanese macaque, reconstructed based on an anatomical model-matching method. J. Hum. Evol. 58, 252–261.

Okada, M., 1985. Primate bipedal walking: Comparative kinematics. In: Kondo, S. (Ed.), Primate Morphophysiology, Locomotor Analyses, and Human Bipedalism. University of Tokyo Press, Tokyo, pp. 47–58.

O’Neill, M.C., Lee, L-F., Larson, S.G., Demes, B., Stern, J.T., Jr., Umberger, B.R., 2013. A three-dimensional musculoskeletal model of the chimpanzee (*Pan troglodytes*) pelvis and hind limb. J. Exp. Biol. 216, 3709–3723.

O’Neill, M.C., Lee, L-F., Demes, B., Thompson, N.E., Larson, S.G., Stern, J.T., Jr., Umberger, B.R., 2015. Three-dimensional kinematics of the pelvis and hind limbs in chimpanzee (*Pan troglodytes*) and human bipedal walking. J. Hum. Evol. 86, 32–42.

O’Neill, M.C., Umberger, B.R., Holowka, N.B., Larson, S.G., Reiser, P.J., 2017. Chimpanzee super strength and human skeletal muscle evolution. Proc. Nat. Acad. Sci. USA 114, 7343–7348.

O’Neill, M.C., Demes, B., Thompson, N.E., Umberger, B.R., 2018. Three-dimensional kinematics and the origin of the hominin walking stride. J. R. Soc. Interface 15, 20180205.

Pilbeam, D.R., Lieberman, D.E., 2017. Reconstructing the last common ancestor of chimpanzees and humans. In: Muller, M.N., Wrangham, R.W., Pilbeam, D.R. (Eds.), Chimpanzees and Human Evolution. Harvard University Press, Cambridge, pp 22–141.

Pontzer, H., Raichlen, D.A., Sockol, M.D., 2009. The metabolic cost of walking in humans, chimpanzees and early hominins. J Hum. Evol. 56, 43–54.

Pontzer, H., Raichlen, D.A., Rodman, P.S., 2014. Bipedal and quadrupedal locomotion in chimpanzees. J. Hum. Evol. 66, 64–82.

Prang, C.T., 2015. Calcaneal robusticity in Plio-Pleistocene hominins: Implications for locomotor diversity and phylogeny. J. Hum. Evol. 80, 135–146.

Prang, C.T., 2019. The African ape-like foot of *Ardipithecus ramidus* and its implications for the origin of bipedalism. eLife 8, e44433.

Raichlen, D.A., Gordon, A.D., 2017. Interpretation of footprints from Site S confirms human-like bipedal biomechanics in Laetoli hominins. J. Hum. Evol. 107, 134–138.

Richmond, B.G., Jungers, W.L., 2008. *Orrorin tugenensis* femoral morphology and the evolution of hominin bipedalism. Science 319, 1662–1664.

Roberts, T.J., 2016. Contribution of elastic tissues to the mechanics and energetics of muscle function during movement. J. Exp. Biol. 219, 266–275.

Roberts, T.J., Chen, M.S., Taylor, C.R., 1998. Energetics of bipedal running. II: Limb design and running mechanics. J. Exp. Biol. 201, 2753–2762.

Rodman, P.S., McHenry, H.M., 1980. Bioenergetics and the origin of human bipedalism. Am. J. Phys. Anthropol. 52, 103–106.

Ross, C.F., Lockwood C.A., Fleagle, J., Jungers, W.L., 2002. Adaptation and behavior in the primate fossil record. In: Plavcan J.M., Kay, R.F., Jungers, W.L., van Schaik, C.P., (Eds.), Reconstructing Behavior in the Primate Fossil Record. Kluwer Academic, New York, pp. 1–41.

Royer, T.D., Martin, P.E., 2005. Manipulations of leg mass and moment of inertia: Effects on energy cost of walking. Med. Sci. Sports Exerc. 37, 646–656.

Ruff, C.B., 1995. Biomechanics of the hip and birth in early *Homo*. Am. J. Phys. Anthropol. 98, 527–574.

Sasaki, K., Neptune, R.R., Kautz, S.A., 2009. The relationships between muscle, external, internal and joint mechanical work during normal walking. J. Exp. Biol. 212, 738–744.

Schache, A.G., Brown, N.A.T., Pandy, M.G., 2015. Modulation of work and power by the human lower-limb joints with increasing steady-state locomotion speed. J. Exp. Biol. 218, 2472–2481.

Schmitt, D., 2003. Insights into the evolution of human bipedalism from experimental studies of humans and other primates. J. Exp. Biol. 206, 1437–1448.

Schmitt, D., Larson, S.G., 1995. Heel contact as a function of substrate type and speed in primates. Am. J. Phys. Anthropol. 96, 39–50.

Schmitz, A., Silder, A., Heiderscheit, B., Mahoney, J., Thelen, D.G., 2009. Differences in lower-extremity muscular activation during walking between healthy older and young adults. J. Electromyogr. Kinesiol. 19, 1085–1091.

Schultz, A.H., 1930. The skeleton of the trunks and limbs of higher primates. Hum. Biol. 2, 303–438.

Schultz, A.H., 1961. Vertebral column and thorax. Primatologia 4,1–66.

Simpson, S.W., Levin, N.E., Quade, J., Rodgers, M.J., Semaw, S., 2019. *Ardipithecus ramidus* postcrania from the Gona Project area, Afar Regional State, Ethiopia. J. Hum. Evol. 129, 1–45.

Slider, A., Whittington, B., Heiderscheit, B., Thelen, D.G., 2007. Identification of passive elastic joint moment-angle relationships in the lower extremity. J. Biomech. 40, 2628–2635.

Sockol, M.D., Raichlen, D.A., Pontzer, H., 2007. Chimpanzee locomotor energetics and the origin of human bipedalism. Proc. Natl. Acad. Sci. USA 104, 12265–12269.

Stansfield, B.W., Nicol, A.C., Paul, J.P., Kelly, I.G., Graichen, F., Bergmann, G., 2003a. Direct comparison of calculated hip joint contact forces with those measured using instrumented implants. An evaluation of a three-dimensional mathematical model of the lower limb. J. Biomech. 36, 929–936.

Stansfield, B.W., Hillman, S.J., Hazlewood, M.E., Lawson, A.M., Mann, A.M., Loudon, I.R., Robb, J.E., 2003b. Normalisation of gait data in children. Gait Posture. 17, 81–87.

Stern, J.T., Jr., Susman, R.L., 1981. Electromyography of the gluteal muscles in *Hylobates*, *Pongo*, and *Pan*: Implications for the evolution of hominid bipedality. Am. J. Phys. Anthropol. 55, 153–166.

Stern, J.T., Jr., Larson, S.G., 1993. Electromyographic study of the obturator muscles in non-human primates: Implications for interpreting the obturator externus groove of the femur. J. Hum. Evol. 24, 403–427.

Susman, R.L., 1983. Evolution of the human foot: Evidence from Plio-Pleistocene hominids. Foot Ankle 3, 365–376.

Susman, R.L., Stern, J.T., Jr., Jungers, W.L., 1984. Arboreality and bipedality in the Hadar hominids. Folia Primatol. 43, 113–156.

Sutherland, D., 1997. The development of mature gait. Gait Posture. 6, 163–170.

Takanashi, K.Z., Worster, K., Bruening, D.A., 2017. Energy neutral: The human foot and ankle subsections combine to produce near zero net mechanical work during walking. Sci. Rep. 7, 15404.

Thorpe, S.K.S., Crompton, R.H., Gunther, M.M., Ker, R.F., Alexander, R.McN., 1999. Dimensions and moment arms of the hind- and forelimb muscles of common chimpanzees (*Pan troglodytes*). Am. J. Phys. Anthropol. 110, 179–199.

Umberger, B.R., Rubenson, J., 2011. Understanding muscle energetics in locomotion: New modeling and experimental approaches. Exerc. Sports Sci. Rev. 39, 59–67.

Vereecke, E., D’Août, K., De Clercq, D., van Elsacker, L., Aerts, P., 2003. Dynamic plantar pressure distribution during terrestrial locomotion of bonobos (*Pan paniscus*). Am. J. Phys. Anthropol. 120, 373–383.

Vrahas, M.S., Brand, R.A., Brown, T.D., Andrews, J.G., 1990. Contribution of passive tissues to the intersegmental moments at the hip. J. Biomech. 23, 357–62.

Ward, C.V., 1993. Torso morphology and locomotion in *Proconsul nyanzae*. Am. J. Phys. Anthropol. 92, 291–328.

Ward, C.V., Hammond, A.S., Plavcan, J.M., Begun, D.R., 2019. A late Miocene hominoid partial pelvis from Hungary. J. Hum. Evol. 136, 102645.

Warrener, A.G., 2017. Hominin hip biomechanics: Changing perspectives. Anat. Rec. 300, 932–945.

Webber, J.T., Raichlen, D.A., 2016. The role of plantigrady and heel-strike in the mechanics and energetics of human walking with implications for the evolution of the human foot. J. Exp. Biol. 219, 3729–3737.

Whittington, B., Slider, A., Heiderscheit, B., Thelen, D.G., 2008. The contribution of passive-elastic mechanisms to lower extremity joint kinetics during human walking. Gait Posture 27, 628–634.

Winter, D.A, Yack, H.J., 1987. EMG profiles during normal human walking: Stride-to-stride and inter-subject variability. Electro. Clinical Neurophys. 67, 402–411.

Woledge, R.C., Curtin, N.A., Homsher, E., 1985. Energetic Aspects of Muscle Contraction. Academic Press, London.

Yoon, Y.S., Mansour, J.M., 1982. The passive elastic moment at the hip. J. Biomech. 15, 905–10.

Yamazaki, N., Ishida, H., Kimura, T., Okada, M., 1979. Biomechanical analysis of primate bipedal walking by computer simulation. J. Hum. Evol. 8, 337–349.

Yamazaki, N., 1985 Primate bipedal walking: Computer simulation. In: Kondo, S. (Ed.), Primate Morphophysiology, Locomotor Analyses, and Human Bipedalism. U Tokyo Press, Tokyo, pp. 105–130.

Zelik, K.E., Kuo, A.D., 2010. Human walking isn’t all hard work: Evidence of soft tissue contributions to energy dissipation and return. J. Exp. Biol. 213, 4257–4264.

Zelik, K.E., Takahashi, K.Z., Sawicki GS. 2015. Six degree-of-freedom analysis of hip, knee, ankle and foot provides updated understanding of biomechanical work during human walking. J. Exp. Biol. 218, 876–886.

Zelik, K.E., Franz, J.R. 2017. It’s positive to be negative: Achilles tendon work loops during human locomotion. PLoS One. 12, e0179976.

Zelik, K.E., Honert, E.C., 2018. Ankle and foot power in gait analysis: Implications for science, technology and clinical assessment. J. Biomech. 75, 1–12.

Zihlman, A., Bolter, D., Boesch, C., 2004. Wild chimpanzee dentition and its implications for assessing life history in immature hominin fossils. Proc. Natl. Acad. Sci. USA 101, 10541–10543.

