## Supplementary Online Materials for "Adaptations for bipedal walking: Musculoskeletal structure and three-dimensional joint mechanics of humans and bipedal chimpanzees (*Pan troglodytes*)"

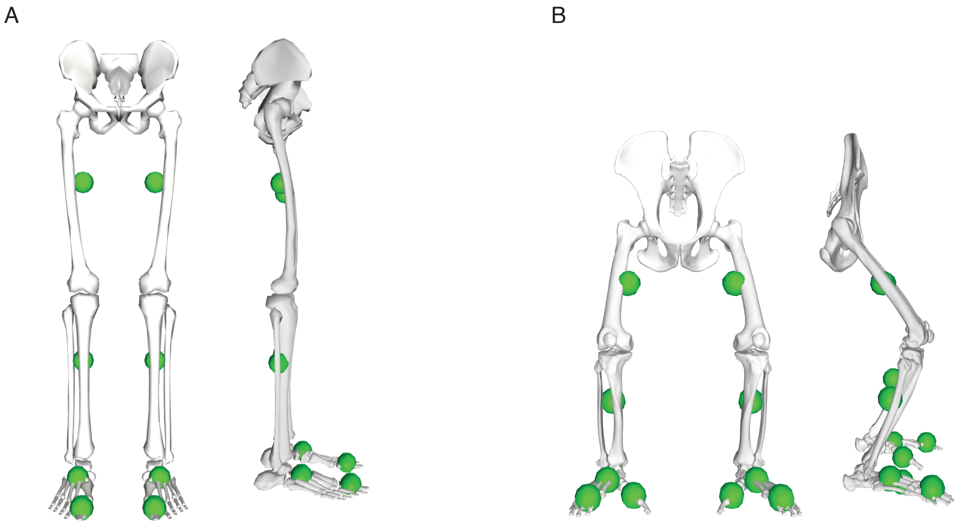


**SOM Figure S1.** Front and side views of the generic A) human (Delp et al., 1990) and B) chimpanzee (O'Neill et al., 2013) pelvis and lower/hind limbs models showing 3-D segment center of mass positions. The generic models were scaled to each human and chimpanzee subject based on the segment lengths and body mass for each experiment. Segment mass 3-D positions are shown as green spheres. Sphere diameter is consistent for all segments and does not reflect mass magnitude.

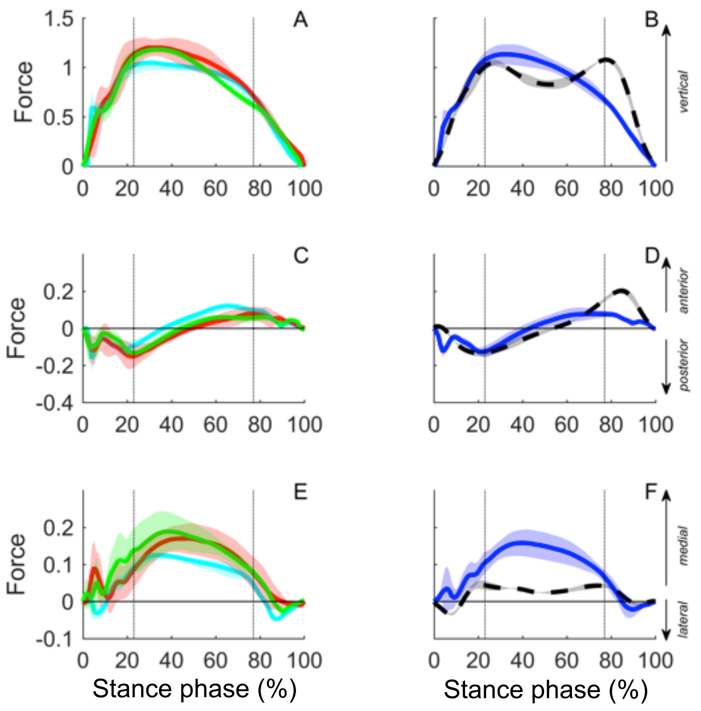


**SOM Figure S2.** The A, B) vertical, C, D) fore-aft and E, F) medio-lateral ground reaction forces over a walking step (mean ± s.d.) for the three chimpanzees (column 1) as well as chimpanzees (solid blue line) and humans (dashed black line) as groups (column 2) at similar dimensional speeds (i.e., absolute speed match: $v = 1.09$ m $s^{-1}$). Forces are in dimensionless units. Stance phase events as in Figure 2.

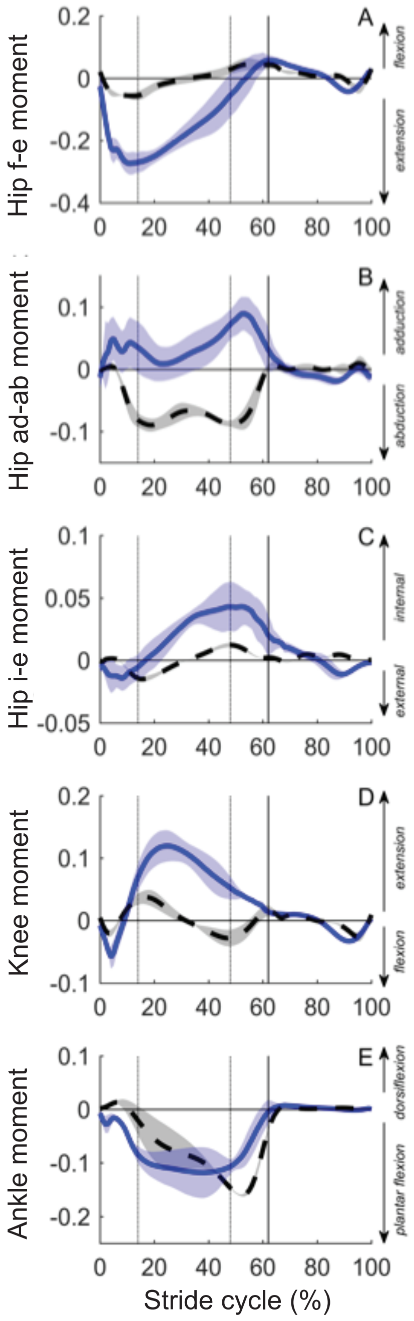


**SOM Figure S3.** The A) hip flexion-extension (f-e) moment, B) hip adduction-abduction (ad-ab) moment, C) internal-external (i-e) rotation, D) knee flexion-extension moment, and E) ankle plantar flexion-dorsiflexion moment over a walking stride (mean ± s.d.) for humans (dashed black line) and chimpanzees (solid blue line) at similar dimensional speeds (i.e., absolute speed match: $v = 1.09$ m s$^{-1}$). Moments are in dimensionless units. Stride cycle events as in Figure 3.

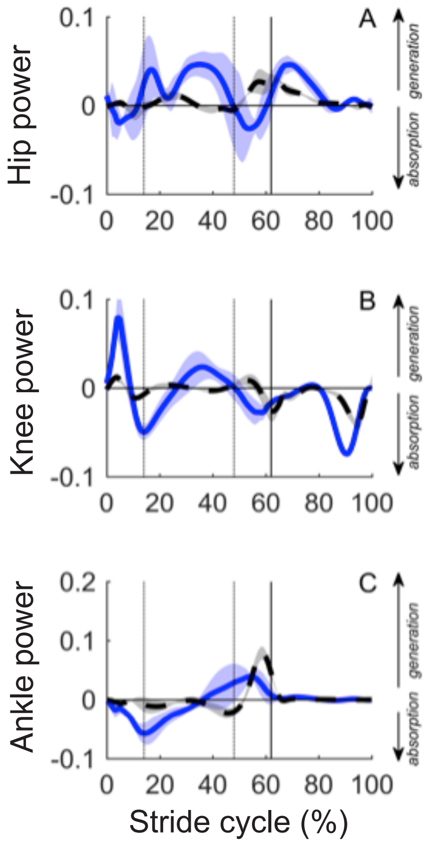


**SOM Figure S4.** The A) 3-D hip power, B) knee flexion-extension power, and C) ankle plantar flexion-dorsiflexion power over a walking stride (mean ± s.d.) for humans (dashed black line) and chimpanzees (solid blue line) at similar dimensional speeds (i.e., absolute speed match: $v$ = 1.09 m $s^{-1}$). Total 3-D joint powers were computed after applying a matrix rotation prior to differentiation of the 3-D angular velocity at the hip. This transformation ensured that the second derivative of the joint angular velocities was equal to joint angular accelerations, and permits the correct instantaneous total power output. Powers are in dimensionless units. Stride cycle events as in Figure 4.

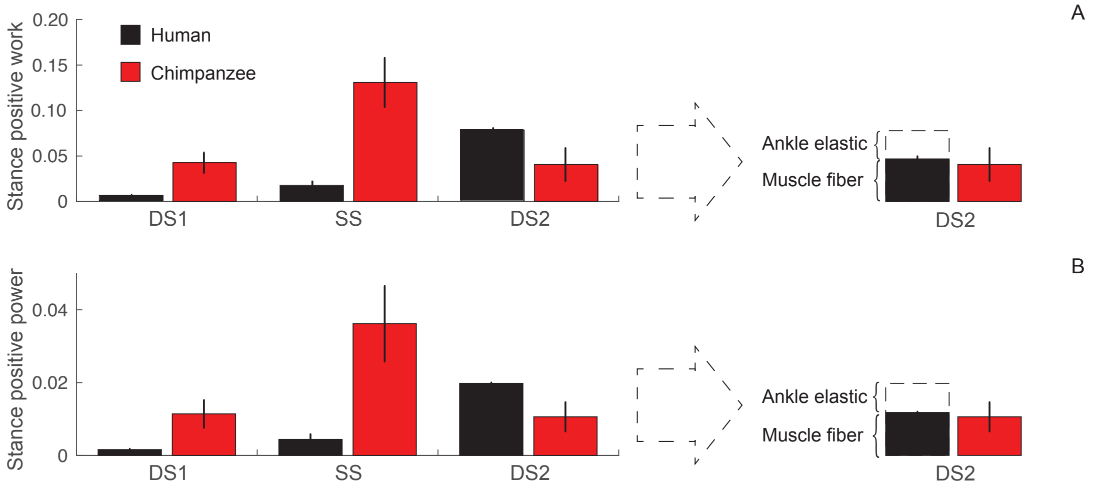


**SOM Figure S5.** The A) stance phase positive joint work and B) power output in the first double support (DS1), single support (SS), and second double-support (DS2) period of a walking stride (mean ± s.d.) at similar dimensional speeds (i.e., absolute speed match: $v$ = 1.09 m $s^{-1}$). The elastic contribution from the ankle to the second-double support (DS2) period. The remaining total mechanical work and power is attributed to 'muscle fiber' mechanics.

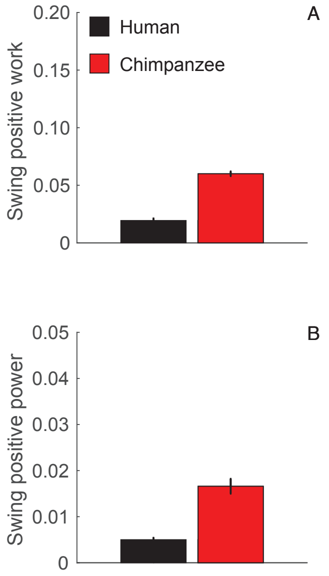


**SOM Figure S6.** The A) swing phase positive joint work and B) power for humans and chimpanzees at similar dimensional speeds (i.e., absolute speed match: $v = 1.09\ \text{m s}^{-1}$).

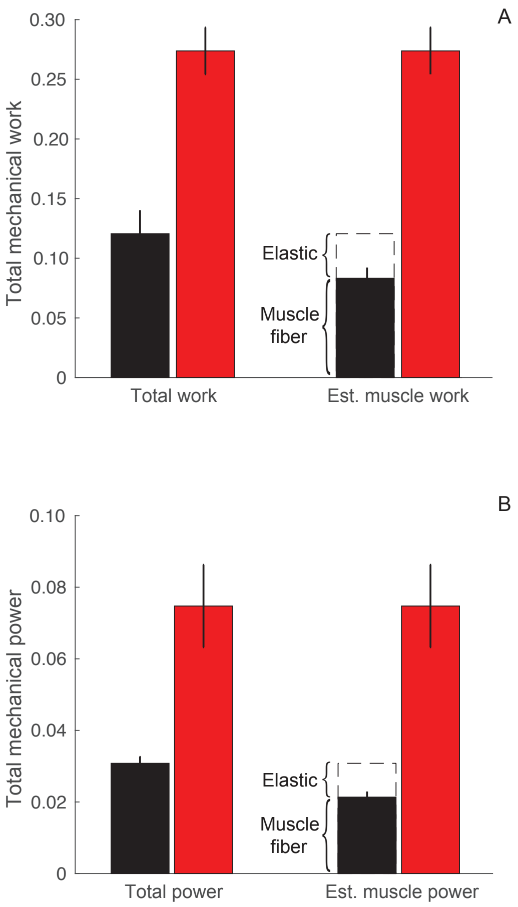


**SOM Figure S7.** The A) total positive limb work and B) average positive limb power over a full stride during walking in humans (black) and chimpanzees (red) (mean ± s.d.) at similar dimensional speeds (i.e., absolute speed match: $v = 1.09$ m s$^{-1}$).

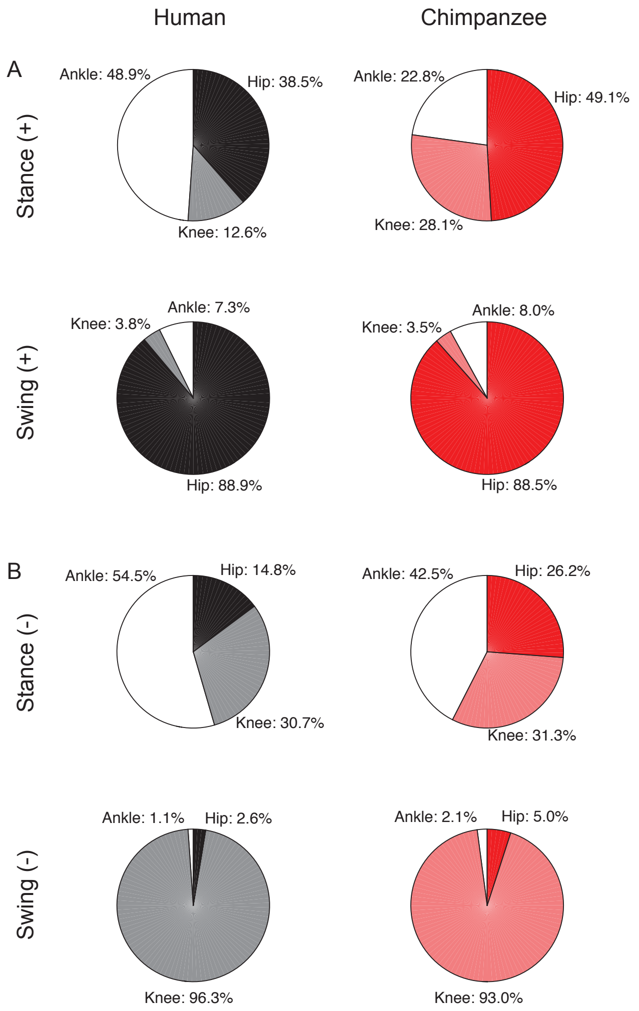


**SOM Figure S8.** The distribution of A) positive mechanical work and B) negative mechanical work among the limb joints during stance and swing in humans (column 1) chimpanzees (column 2) at similar dimensional speeds (i.e., absolute speed match: $v$ = 1.09 m $s^{-1}$).

**SOM Table S1**

Segment mass, length and inertial properties of the generic chimpanzee (O'Neill et al., 2013) and human (Delp et al., 1990) models. Human parameters were taken from Delp et al. (1990). For chimpanzees, body mass, segment lengths, and diameters were measured on each chimpanzee subject 2 to 4 times during the data collection period following published protocols (e.g., Raichlen, 2004; Schooneart et al. 2007). Ellipsoid models were used to derive segment masses, moments of inertia, and center of mass (COM) positions (Raichlen, 2004). The dimensionless parameters were averaged within and then across subjects, with geometric scaling then used to derive parameters for the generic chimpanzee model. The generic chimpanzee (mass, $M_b = 55.0$ kg) and human (mass, $M_b =$ 75.2 kg) model parameters were then scaled to each subject prior to inverse dynamics calculations (c.f., O'Neill et al., 2015).

| Segment | Segment mass[a] (kg) | | Segment length[b] (m) | | Segment moment of inertia[c] (kg $m^{-2}$) | | COM position[c] (m) | |
|---|---|---|---|---|---|---|---|---|
| | Chimpanzee | Human | Chimpanzee | Human | Chimpanzee | Human | Chimpanzee | Human |
| Thigh | 4.125 | 9.3014 | 0.265 | 0.396 | xx: 0.0224<br>yy: 0.0059<br>zz: 0.0220 | xx: 0.1339<br>yy: 0.0351<br>zz: 0.1412 | x: 0.000<br>y: –0.140<br>z: 0.008 | x: 0.0000<br>y: –0.1700<br>z: 0.0000 |
| Leg | 1.705 | 3.7075 | 0.280 | 0.429 | xx: 0.0064<br>yy: 0.0011<br>zz: 0.0062 | xx: 0.0504<br>yy: 0.0051<br>zz: 0.0511 | x: –0.030<br>y: –0.130<br>z: –0.005 | x: 0.0000<br>y: –0.1867<br>z: 0.0000 |
| Foot | 0.900 | 1.350 | 0.115 | 0.139 | xx: 0.003<br>yy: 0.004<br>zz: 0.004 | xx: 0.0014<br>yy: 0.0039<br>zz: 0.0041 | x: 0.050<br>y: –0.020<br>z: 0.000 | x: 0.1000<br>y: 0.0300<br>z: 0.0000 |
| Toes | 0.130 | 0.2166 | 0.079 | 0.071 | xx: 0.001<br>yy: 0.001<br>zz: 0.001 | xx: 0.0001<br>yy: 0.0002<br>zz: 0.0001 | x: 0.025<br>y: 0.000<br>z: 0.000 | x: 0.0346<br>y: 0.0060<br>z: –0.0175 |
| Hallux | 0.050 | — | 0.105 | — | xx: 0.001<br>yy: 0.001<br>zz: 0.001 | — | x: 0.020<br>y: –0.015<br>z: –0.030 | — |

[a] Human foot mass includes model talus + calcaneus segments from Delp et al. (1990). Toe masses includes digits 1–5 (human) or digits 2–5 (chimp), with a separate digit 1 mass for the chimp model.
[b] Segment lengths are measured from joint coordinate centroids of each segment with the model in neutral position (c.f. de Leva, 1996). 'Foot length' is measured from the ankle joint centroid to metatarsophalangeal joint centroid for digits 1–5 (human) or 2–5 (chimp). Hallux length is measured from the tarsometatarsal joint centroid to the tip of digit 1.
[c] *xx*, *yy*, *zz* and *x, y, z* are given with respect to the segment's local reference frame.

**SOM Table S2**

Spatiotemporal gait parameters in human walking. Human data are matched to the chimpanzee data in dimensional (i.e., absolute-speed match) form (mean ± s.d.).

| | $v$[a] (m $s^{-1}$) | $v$[b] | $Fr$[c] | Stance time (s) | Swing time (s) | Stride length (m) | Stride frequency (Hz) | Duty factor |
|---|---|---|---|---|---|---|---|---|
| Humans | 1.08 ± 0.02 | 0.36 ± 0.01 | 0.13 ± 0.01 | 0.79 ± 0.05 | 0.42 ± 0.03 | 1.31 ± 0.07 | 0.83 ± 0.05 | 0.66 ± 0.00 |

[a] Dimensional velocity.
[b] Dimensionless velocity.
[c] Froude number.

**SOM Table S3**

Adjusted coefficient of multiple correlation of ground reaction forces ($F_y$, $F_x$, $F_z$) in dimensional (i.e., absolute-speed match) form (mean ± s.d.).

| | $F_{y,\ vert}$ | $F_{x,\ ant\text{-}post}$ | $F_{z,\ med\text{-}lat}$ |
|---|---|---|---|
| Humans[a] | 0.994 ± 0.003 | 0.996 ± 0.001 | 0.965 ± 0.017 |
| Among humans[b] | 0.958 ± 0.018 | 0.972 ± 0.007 | 0.916 ± 0.038 |

Human data are matched to the chimpanzee data in dimensionless (i.e., relative-speed match) form (mean ± s.d.). Coefficient of multiple correlation mean ± s.d. for 4 trials per subject, $n = 3$ subjects. The y, x, z components of the ground reaction forces are vertical (vert), anterior-posterior (ant-post) and medial-lateral (med-lat).

[a] Correlations between strides within subjects.

[b] Correlations between subjects.

**SOM Table S4**

Peak joint moments in humans in dimensionless units (i.e., absolute-speed match) form (mean ± s.d.).

| | Hip | | | Knee | Ankle |
|---|---|---|---|---|---|
| | Flexion-extension | Adduction-abduction | Internal-external rotation | Flexion-extension | Plantar flexion-dorsiflexion |
| Humans | –0.061 ± 0.009 (ext.) | –0.093 ± 0.009 (abd.) | 0.016 ± 0.002 (int.) | 0.041 ± 0.009 (ext.) | –0.161 ± 0.004 (plantar) |

The direction of the peak joint moment is given as flexion (flex.), extension (ext.), abduction (abd.), adduction (add.), internal rotation (int.), external rotation (ext.), plantar flexion (plantar.) or dorsiflexion (dorsi.).

**SOM Table S5**

Peak joint power in humans in dimensionless units (i.e., absolute-speed match) form (mean ± s.d.).

| | Hip power | | Knee power | | Ankle power | |
|---|---|---|---|---|---|---|
| | + | – | + | – | + | – |
| Humans | 0.033 ± 0.009 | –0.011 ± 0.006 | 0.016 ± 0.002 | –0.042 ± 0.006 | 0.080 ± 0.010 | –0.023 ± 0.009 |

**SOM References**

de Leva, P., 1996. Joint center longitudinal positions computed from a selected subset of Chandler's data. J. Biomech. 29, 1231–1233.

Delp, S.L., Loan, J.P., Hoy, M.G., Zajac, F.E., Topp, E.L., Rosen, J.M., 1990. An interactive graphics-based model of the lower extremity to study orthopaedic surgical procedures. IEEE Trans. Biomed. Eng. 37, 757–767.

O'Neill, M.C., Lee, L.F., Larson, S.G., Demes, B., Stern, J.T. Jr., Umberger, B.R., 2013 A three-dimensional musculoskeletal model of the chimpanzee (*Pan troglodytes*) pelvis and hind limb. J. Exp. Biol. 216, 3709–3723.

O'Neill, M.C., Lee, L.F., Larson, S.G., Demes, B., Thompson, N.E., Stern, J.T. Jr., Umberger, B.R., 2015. Three-dimensional kinematics of the pelvis and hind limbs in chimpanzee (*Pan troglodytes*) and human bipedal walking. J. Hum. Evol. 86, 32–42.

Raichlen, D.A., 2004. Convergence of forelimb and hindlimb natural pendular periods in baboons (*Papio cynocephalus*) and its implication for the evolution of primate quadrupedalism. J. Hum. Evol. 46,719–738.

Schoonaert, K., D'Août, K., Aerts, P., 2007. Morphometrics and inertial properties in the body segments of chimpanzees (*Pan troglodytes*). J. Anat. 201, 518–531.
